# CRISPR-TRAPSeq identifies the QKI RNA binding protein as important for astrocytic maturation and control of thalamocortical synapses

**DOI:** 10.1101/2020.03.13.991224

**Authors:** Kristina Sakers, Yating Liu, Lorida Llaci, Michael J. Vasek, Michael A. Rieger, Sean Brophy, Eric Tycksen, Renate Lewis, Susan E. Maloney, Joseph D. Dougherty

## Abstract

Quaking RNA binding protein(QKI) is essential for oligodendrocyte development as myelination requires MBP mRNA regulation and localization by the cytoplasmic isoforms(e.g. QKI-6). QKI-6 is also highly expressed in astrocytes, which were recently demonstrated to have regulated mRNA localization. Here, we show via CLIPseq that QKI-6 binds 3’ UTRs of a subset of astrocytic mRNAs, including many enriched in peripheral processes. Binding is enriched near stop codons, which is mediated partially by QKI binding motifs(QBMs) yet spreads to adjacent sequences. We developed CRISPR TRAPseq: a viral approach for mosaic, cell-type specific gene mutation with simultaneous translational profiling. This enabled study of QKI-deleted astrocytes in an otherwise normal brain. Astrocyte-targeted QKI deletion altered translation and maturation, while also increasing synaptic density within the astrocyte’s territory. Overall, our data indicate QKI is required for astrocyte maturation and demonstrate an approach for a highly targeted translational assessment of gene knockout in specific cell-types *in vivo.*

## Introduction

Post-transcriptional regulation of mRNA by RNA binding proteins(RBPs) is pervasive in the Central Nervous System(CNS). RBPs serve as the *trans*-acting factors that recognize *cis*-acting elements that are commonly found in the 3’ UTR of localized transcripts^1^. These proteins may directly transport RNAs to distal processes^2^, alter transcript stability, or control translation^3,4^, for example to prevent ectopic expression during mRNA transport, or to activate translation in response to a molecular cue. While these processes have been widely studied in other cell-types, it has recently become apparent that astrocytes also must have substantial cytoplasmic posttranscriptional regulation: for example, a set of mRNAs enriched on ribosomes in peripheral astrocyte processes(PAPs) have been defined, consistent with sequence-regulated local translation^5^. Likewise, there are clear examples of posttranscriptional regulation, via expression of specific cytoplasmic RBPs, being an essential aspect of differentiation and maturation of neurons^6^, microglia^7^, and oligodendrocytes^8^. However, aside from elegant work on the Fragile X Mental Retardation Protein(FMRP)^9,10^, post-transcriptional regulators of such processes in astrocytes largely remain unknown.

QKI is the mammalian homolog of *C. elegans* GLD-1 and *D. Melanogaster* HOW, and part of the signal transduction and activation of RNA(STAR) family of post-transcriptional RBPs. Members of this evolutionarily conserved family of proteins are characterized by conserved domains which are necessary for homodimerization and RNA binding^11^. STAR family proteins are important early in development as their expression regulates smooth muscle development, cell-fate, and germ-line development^12,13^. However, QKI expression in mice remains high in brain throughout the lifespan^14,15^, suggesting ongoing regulatory roles in the CNS. QKI is predominantly expressed in glia in the brain^16^ and its function in oligodendrocytes has been well-studied: QKI plays a role in oligodendrocyte differentiation and development by regulating the stability of mRNAs that inhibit cell cycle progression and promote differentiation^17–19^. Further, cytoplasmic isoforms of QKI control the nuclear export, stability, and localization of myelin basic protein(MBP) mRNA to oligodendrocyte processes^8,20^.

Although most of our knowledge of QKI’s function *in vivo* comes from studies focused on oligodendrocytes, this protein is also expressed in astrocytes in both mice and humans^21,22^. In an astroglioma cell line, it has been shown that one QKI isoform stabilizes transcripts, particularly those induced by interferons^23^. Moreover, examination of PAP localized transcripts from maturing astrocytes revealed a preponderance of QKI-binding motifs(QBMs) in their UTRs^5^. However, we still do not understand to which cytoplasmic targets QKI binds in vivo nor what the downstream effects on these targets are in the maturing postnatal brain. Furthermore, study of this gene is complicated as full knockouts are embryonic lethal^12^, making it challenging to disentangle direct cellular function of QKI from indirect effects of disrupted development. Here, we focus on the cytoplasmic isoform QKI-6 to comprehensively define its targets in the maturing brain. We test the hypothesis that QKI-6 disproportionately binds PAP-localized transcripts and define the role of QKI in mRNA ribosome occupancy for astrocytes *in vivo*. We find, by Cross-Linking and Immunoprecipitation(CLIP)-seq, many QKI-6 targets are indeed astrocytic and overlap with transcripts enriched on PAP ribosomes. Further, we develop a new approach to define the role for this gene on ribosome occupancy in astrocytes *in vivo*. We demonstrate that loss of QKI affects ribosome occupancy and maturation, and finally that this has ultimate consequences on synapse numbers.

## Results

### QKI-*6 protein binds translationally regulated mRNAs in astrocytes*

To understand how QKI regulation of mRNAs might affect astrocyte development and/or function, we first sought to determine which mRNAs were targeted by QKI *in vivo*. Although a few targets of QKI have been well studied in oligodendrocytes^18,20,24^, and to a lesser extent in astrocytes^25,26^, a compendium of all *in vivo* postnatal QKI targets has yet to be compiled. QKI has three major isoforms in humans and in mice, designated by their unique C-termini^27^, all of which are present in astrocytes^16^(**Fig.1A**) as shown by colocalization with RPL10a-eGFP in Aldh1L transgenic bacTRAP mice. We chose to focus on QKI-6 for CLIP experiments for two reasons: firstly, QKI-5 is predominantly nuclear but QKI-6 and QKI-7 are both nuclear and cytoplasmic, and we posited that cytoplasmic isoforms are more likely candidates for translation regulation. Secondly, QKI-6 is expressed earlier in development than QKI-7^28^ suggesting an importance in early neuronal circuitry formation, a function largely governed by astrocytes. Finally, QKI-7 was had more variable intensity of expression across astrocytes(**Fig.1A**).

To define QKI-6 targets in all cells of maturing mouse forebrains from independent biological replicates were homogenized and UV irradiated to cross-link(CL) protein-RNA complexes(**Fig.1B**). We chose P21 because we previously found at this age that mRNAs enriched on PAP ribosomes contain more QBMs than expected by chance. Subsequent QKI-6 immunoprecipitation(IP) or IgG IP(background control) were performed on CL lysates. QKI-6 IP clearly yielded more end-labelable RNA than controls, and a shift in QKI-6/mRNA complexes was detected with decreasing RNAse concentrations, as expected(**Fig. S1**). After generation of CLIPseq libraries from QKI-6 CLIP, IgG IP, and Input RNA(to control for starting transcript abundance) we identified genome-wide significant peaks(p < 0.1). We processed the 6,789 most abundant peaks to identify high-confidence QKI targets: we utilized differential expression pipelines to identify the subset of peaks that had significantly more counts in QKI-6 CLIP(fold change > 2, FDR < 0.05), than both IgG and Input samples. This intersectional analysis conservatively identified 437 peaks from 120 unique transcripts that were used in downstream analyses(**Table S1**). These included the well-studied QKI target *Mbp*(**Fig.1C,D**), as well as Carboxypeptidase E (*Cpe*), which contains predicted QKI response elements(QREs): a full QBM (ACUAAY) that is within 1-20 bases of a second QBM half site (YAAY). Further, unbiased motif discovery^29^in all peaks identified a motif that matched QKI’s QBM^30^(**Fig. 1E**). We further confirmed a subset of targets by IP RT-QPCR in independent samples(**Fig. S2A-C**). Thus, we are confident that the mRNAs discovered via CLIP are *bona fide* QKI targets, *in vivo*.

**Figure 1:**
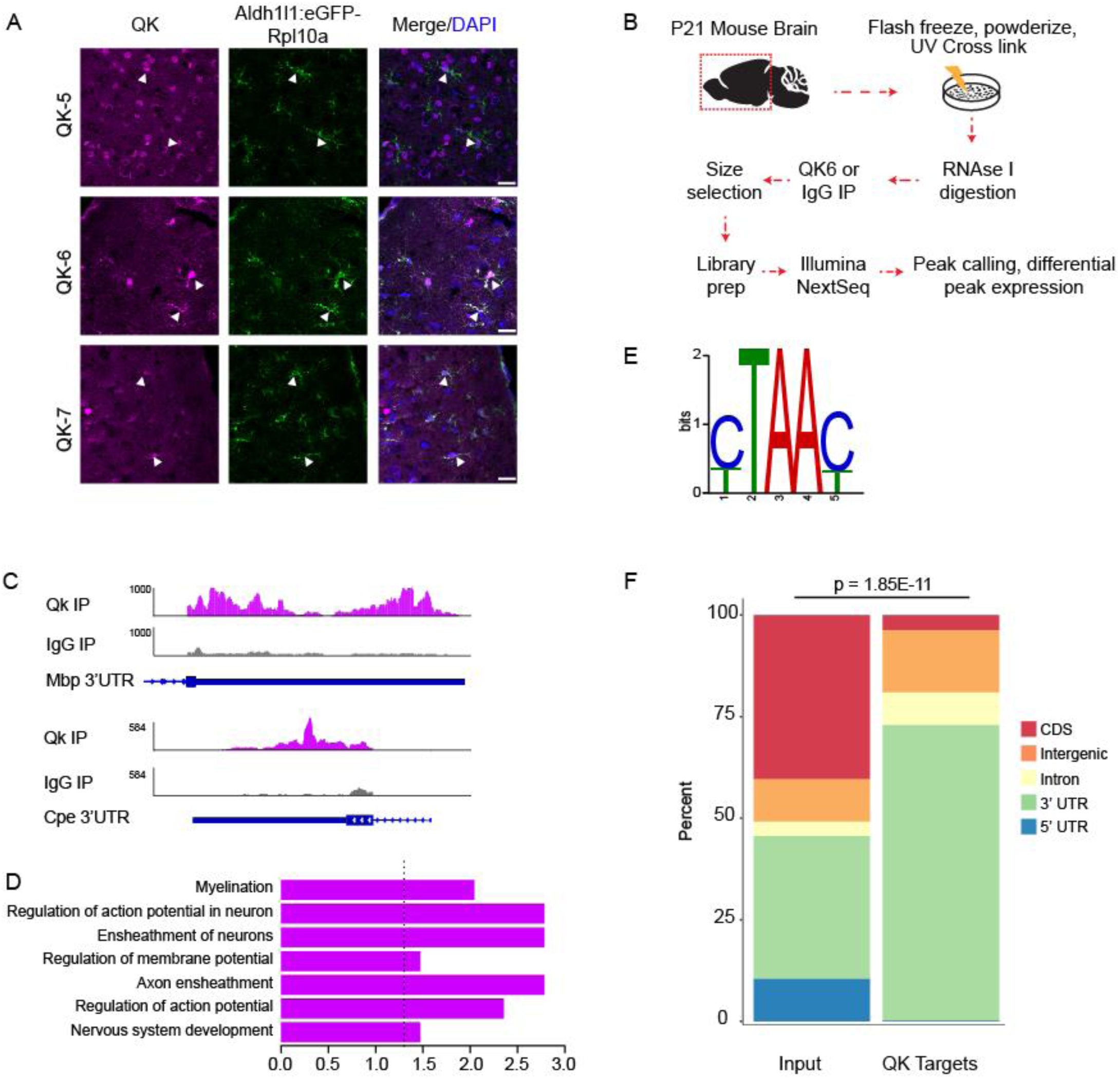
QKI-6 CLIP defines binding targets in developing forebrain *in vivo*. **A)** Representative IHC images reveal all QKI isoforms(magenta) are present in cortical astrocytes, *in vivo*. Scale bar=15μm. **B)** QKI CLIP work flow. **C)** Browser shots of QKI-6 IP vs IgG control reads along Mbp 3’ UTR(top) and Cpe 3’ UTR(bottom) as a control. **D)** Gene Ontology results for all significant biological processes across QKI-6 binding targets. Dotted line represents p = 0.05. **E)** Top identified motif under QKI-6 peaks matches QBM. **F)** Distribution of genome features for significant peaks in QKI-6 IP reveals enrichment of 3’ UTRs.

We next used these data to query the binding properties and target properties of QKI-6. First, we analyzed the genomic distribution of peaks. Generally, RBPs involved in splicing bind highly in introns while those that modulate translation, such as mediating ribosome occupancy, transport and stability of mRNAs, often bind UTRs^31^. We found that the majority(72%) of QKI-6 peaks are in the 3’ UTR, consistent with a potential role in translation regulation, and only a small fraction(8%) were in introns(**Fig. 1F**) suggesting a minimal role of QKI-6 in splicing. These findings are in contrast to a recent pan-QKI CLIP study in cultured myoblasts, in which 38% of the targets are in introns^32^. This large intronic signal is likely due to the splicing and nuclear retention functions of the nuclear isoform, QKI-5^20,32^. Indeed, a recent QKI-5 specific CLIP experiment in E14.5 mouse brain, where QKI-5 is highly expressed in neural stem cells, showed 41.6% of binding sites in introns^28^. Recent work in *C. elegans* emphasized the importance of 5’ UTR QKI motifs^33^, yet we found very few(<1%) QKI-6 peaks in the 5’ UTR of mouse brain transcripts(**Fig. 1F**), consistent with *in vitro* studies that identified only 2% of pan-QKI target peaks are in 5’ UTRs^32^. Thus, with its strong enrichment on 3’ UTRs, QKI-6’s binding pattern appears poised to enable translation regulation rather than splicing.

To better understand potential roles for QKI-6, we then studied the attributes of these mRNAs, and the pattern of binding within their 3’ UTRs. Examining UTR length distributions, it was clear that QKI-bound UTRs are on average longer than randomly selected UTRs from the genome(**Fig. 2A**), suggesting an evolution for greater potential for sequences mediating regulation. To test this, we selected a set of random length-matched UTR sequences for controlsand examined the evolutionary conservation and relative binding positions of QKI. We observed that QKI bound sites were more highly conserved than random UTR sites(**Fig 2B,C).** Examining the pattern of binding within the UTR, there seemed to be a enrichment near the stop codon, with a pattern towards peaks flanking it on either side, as well as a tendency for binding 250 bp upstream of PolyA signal(PAS) sequences. However, there was no difference in the number of PASs between QKI targets and controls, suggesting QKI does not have a general role in alternative polyadenylation(**Fig 2D)**. In length-normalized analyses, there were peaks near both the PAS and the stop codon (**Fig. 2E-G)**, suggesting some interaction with both features. Complimentary to our *de novo* motif discovery(**Fig. 1D**), when we examined average read-depth around predicted QBMs in the CLIP targets, binding was clearly enriched at these motifs(**Fig. 2H)**. Interestingly, QKI binding appears to extend from this primary high-affinity interaction to flanking nucleotides, particularly downstream. In addition to having multiple RNA interacting domains in each protein, QKI is known to homodimerize^11,34^. Thus, our observations could be consistent with cooperativity across QKI molecules: initial binding to a high affinity motif and homodimerization might spread binding for hundreds to thousands of nucleotides along the 3’ UTR. This could also enhance folding of the UTR to bring other parts of the mRNA in contact with the same molecule: head-to-head binding of QKI molecules is thought to strongly fold RNA when binding two sites on the same molecule^34^. Finally, examining all peaks, regardless of QBM-presence, revealed a similar pattern of binding spread(**Fig. 2I).**

**Figure 2:**
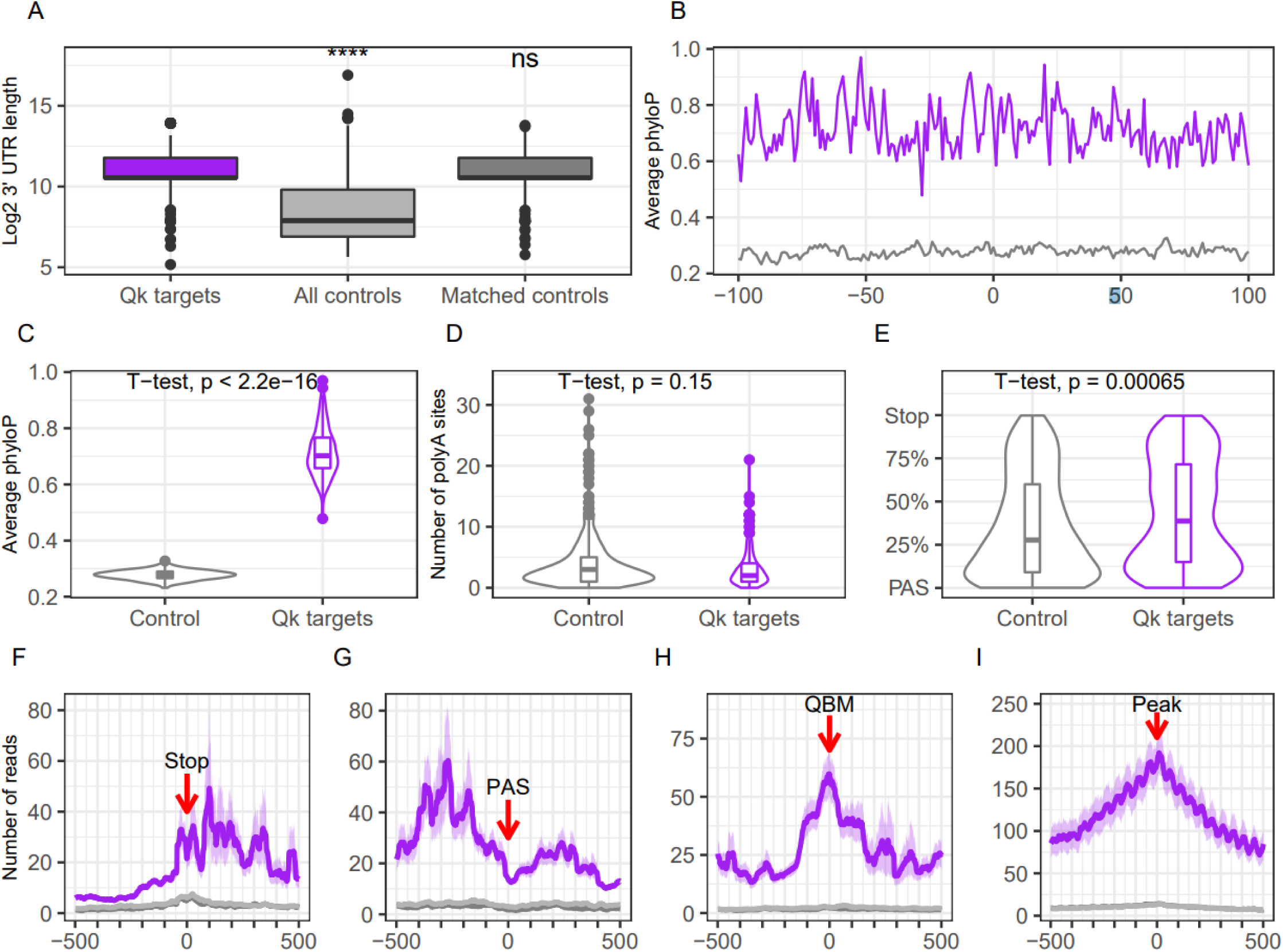
QKI-6 binds conserved UTR regions in a cooperative manner. **A)** UTRs bound by QKI-6 are significantly longer than average UTRs. Rightmost plot: matched-length controls selected for B-G. **B,C)** QKI-6 bound sites are more conserved than random sites in matched-length controls. **D)**QKI-6 peaks are not enriched for polyA sites. **E)** Average of length normalized UTRs show QKI-6 binding closer to stop codon (100%), but with signal near first PAS(0%). **F)** Metagene of QKI-6 shows binding is elevated at stop codon, with flanking peaks, and extends substantially, especially 3’. Purple= QK-6 CLIP, light gray =Input, dark gray =IGG. **G**) Metagene of QKI-6 shows binding is elevated at PAS, with flanking peaks, and extends substantially, especially 5’. QKI-6 binding at any **H)** QBM or **I)** peak extends to flanking sequence.

Next, we sought to examine the role QKI-6 has specifically in astrocytes. QKI is documented as glial-expressed^16^, however its function has primarily been investigated in oligodendrocytes. To determine the relative role in astrocytes, we assessed the percentage of QKI-6 targets highly expressed in different neural cell-types. Using Cell-type Specific Expression Analysis(CSEA)^35^, we mapped the distribution of QKI-6 targets to cells in the brain(**Fig. 3**). Not surprisingly, we found that 34% of targets came from myelinating oligodendrocytes as QKI expression is dramatically upregulated in these cells compared to oligodendrocyte precursors^36^. This is consistent with our analysis of QKI-bound targets using Gene Ontology, where we identified enrichment of myelin-associated transcripts(**Fig. 1D, Table S2**). However, when examining the CSEA data(**Fig. 3B)**, astrocyte-specific transcripts represented an equal percentage of QKI-6 targets as myelinating oligodendrocytes, emphasizing that QKI may play an equally important role in translation regulation in astrocytes.

**Figure 3:**
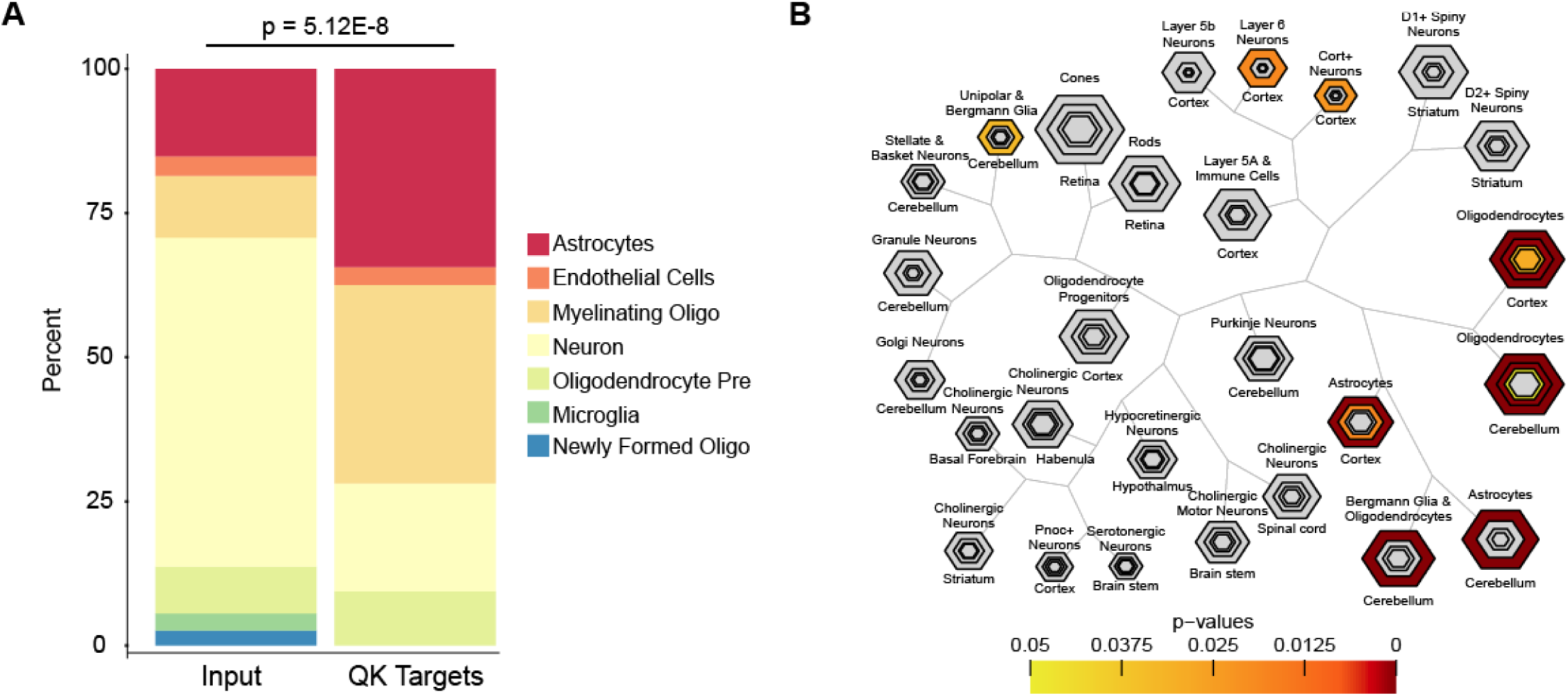
QKI-6 targets membrane protein transcripts in astrocytes. **A)** QKI-6 targets are significantly enriched in transcripts from astrocytes compared to Input transcripts. **B)** Unbiased CSEA analysis^35^ shows QKI-6 targets are most strongly enriched in transcripts enriched in astrocytes and oligodendrocytes. *Hexagons represent list of genes enriched in each cell type going from pSI threshold to include larger but less stringent gene lists (in this cell type and a few others) to thresholds for the most stringent subsets (smallest, central hexagons). Fisher exact testing is used to establish p-value for each CLIP target and each gene list at each threshold and corrected by Benjamini-Hochberg for the number of cell types(color bar). Results consistent across more than one threshold hexagon per cell type are viewed with greater confidence.*

Within astrocytes, we have previously defined a set of transcripts enriched on ribosomes in the peripheral astrocyte process(PAP) and thus represent transcripts likely undergoing translational regulation either to control translation efficiency or localization. To test whether QKI-6 may indeed be playing a role in regulating these transcripts, we compared our CLIP data to our prior analysis. We found that 22% of PAP-enriched mRNAs are also QK CLIP targets(vs 4% of Input genes, Fisher’s Exact Test(FET) p<2.2E-16). Together, these data define QKI-6 targets, *in vivo*, and indicate a significant proportion of these targets are found in 3’ UTRs of astrocyte-expressed transcripts. Furthermore, the significant overlap with PAP-enriched transcripts suggests a role for QKI in regulation of translation of a subset of these transcripts in astrocytes.

### *CRISPR-TRAPseq of* QKI *protein in astrocytes shows regulation of ribosome occupancy of direct targets*

RNA binding proteins can have multiple roles on their targets, including alteration of translation initiation(thus altering ribosome occupancy), mRNA localization, and transcript stability(indirectly impacting levels on ribosomes). As any of these three will have consequences on ribosome occupancy, we asked which QKI-6 binding event might have functional consequences. To address this, we designed an approach to quantify ribosome occupancy by mRNAs after QKI deletion in maturing astrocytes *in vivo*. Because QKI knockouts are not viable^12^, and the most commonly used mouse model(quaking viable, QK^v^) is an enhancer deletion which impacts only oligodendrocytes^16^, we adopted a novel strategy, CRISPR-TRAPseq, to delete QKI in astrocytes and assess changes in ribosome occupancy in these cells. We designed the deletion to be mosaic – impacting a subset of GFAP+ cells in otherwise normally developing brains - to mitigate most non-cell-autonomous effects from mutation. Specifically, we generated knock-in mice containing a loxP-stop-loxP (LSL) Cas9 allele in the Rosa locus that can be expressed under a Cre inducible promoter(**Fig. S3**). We then crossed heterozygotes of this line with mice homozygous for the LSL-Translating Ribosome Affinity Purification(TRAP) construct in the same locus**(Fig.4A)**. We then transcranially injected entire litters of perinatal pups with an AAV expressing Cre and CFP-myc under the control of a GFAP promoter, along with gRNAs targeting QKI under a second promoter(U6), to generate mosaic deletions. Thus, subsequent to GFAP:Cre activity, *all* transduced GFAP positive cells will express the TRAP allele, allowing for cell-type specific enrichment of tagged ribosomes. Further, 50% of these animals will also express Cas9, thus mutating QKI through nuclease cleavage and non-homologous end joining, inducing point mutations and indels into the coding sequence.

First, we confirmed that our Cas9 strategy for gene disruption was effective by immunofluorescence assessing QKI-6 protein(**Fig.4B)**. In P21 animals, we saw robust expression of CFP within cells that exhibited typical astrocyte morphology. Immunofluorescence revealed that cells showed median 79% loss of QKI-6 protein in Cas9+ animals, significantly different from Cas9-littermates(**Fig.4C**). We then conducted TRAPseq on additional animals of each Cas9 genotype. While yields were too low to conduct PAP-TRAP, standard TRAP yielded high quality(RIN>7.7) RNA across replicates, which was then sequenced to a depth of > 27M reads/sample. Cas9 was detectable in the RNAseq, confirming the genotype of each animal. Further, comparison to total RNA ‘Input’ controls reveal a robust enrichment of known astrocyte expressed genes, and depletion of neuronal genes(**Fig. 4D**), indicating the approach was yielding RNA from descendants of perinatal GFAP expressing cells(e.g. astrocytes). Indeed, this relative enrichment of positive control transcripts, and depletion of negative control transcripts was comparable to our prior experiments utilizing our Aldh1L1 bacTRAP animal, validating the efficacy of the method(**Fig S4)**.

**Figure 4:**
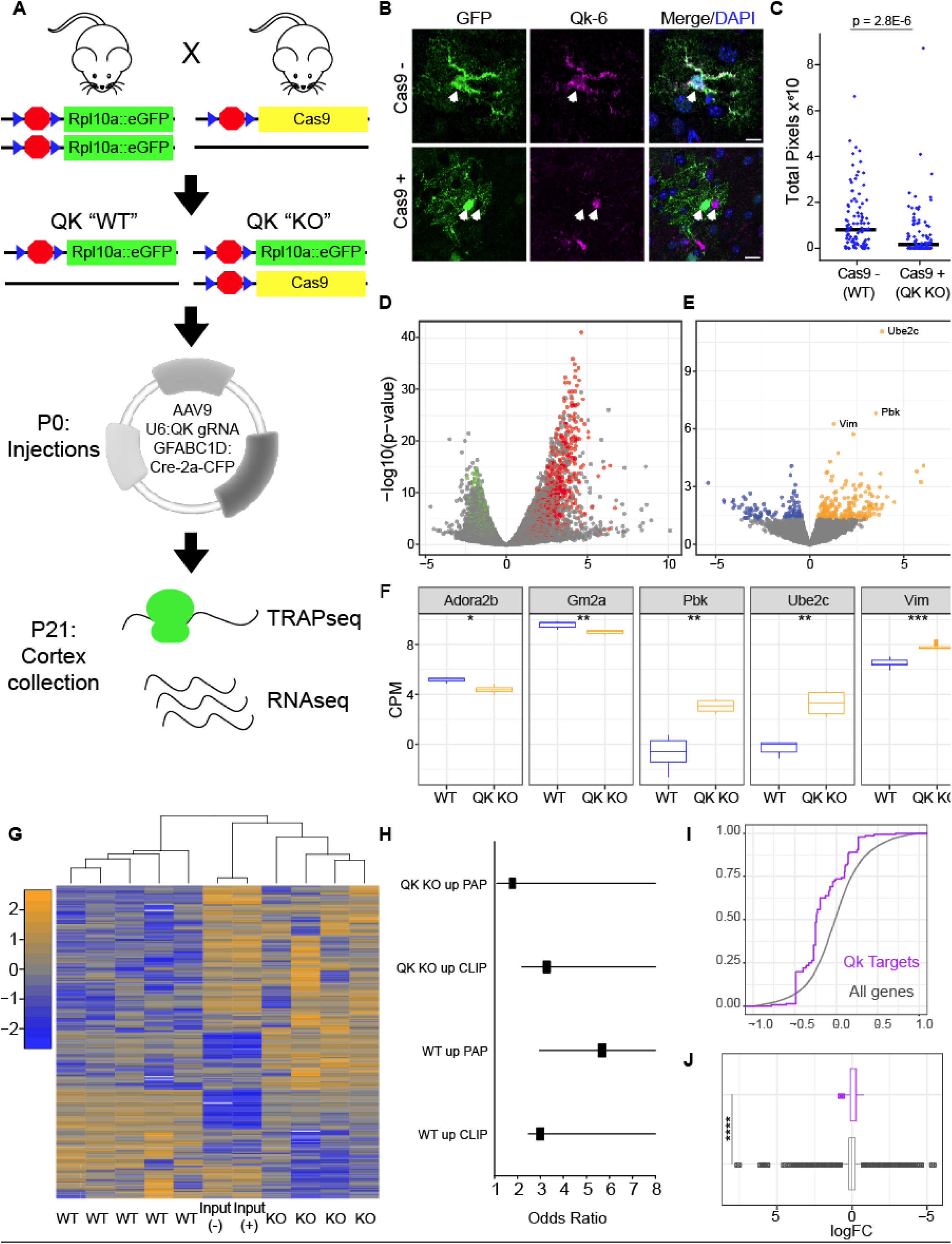
CRISPR-TRAPseq-mediated QKI deletion in maturing astrocytes alters target transcript ribosome occupancy. **A)** Cartoon strategy of QKI knockdown in cortical astrocytes. **B)** Transduced CFP-positive cells(detected by GFP antibody) from GFAP:CFP QKI gRNA-injected mice(Cas9-(top) and Cas9+(bottom)) reveal loss of QKI-6 immunoreactivity in Cas9+ animals, but not in Cas9-animals or in non-transduced adjacent cells. **C)** Quantification of QKI pixels in CFP+ cells across genotypes reveals median 79% reduction in Cas9+ cells. P value represents Wilcoxon rank-sum test. N = 129 cells per genotype **D)** Volcano plot showing mRNA enriched by TRAP from GFAP-Cre gRNA transduced P21 astrocytes is highly enriched in previously defined markers of astrocytes(red) and depleted in markers on neurons(green)^57^, confirming method. **E)** TRAP analysis identifies transcripts with altered ribosome occupancy *in vivo* following QKI mutation. **F)** Boxplots of individual genes with altered ribosome occupancy following QKI deletion. **G)** Heatmap of all TRAP and Input samples showing transcripts with altered ribosome occupancy. **H)** Forest plot illustrates overlap between QKI responsive transcripts, CLIP genes, and locally translated genes as defined^5^. **I)** Cumulative distribution function(CDF) and boxplot (**J**) of CLIP targets expressed in astrocytes show a shift in ribosome occupancy after QKI deletion.

Then, we examined the consequences of QKI mutation on ribosome occupancy by mRNA in astrocytes. Given that the deletion was occurring across a period of postnatal astrocyte maturation, and that the mutation in QKI will impact all isoforms of QKI protein, we expected the resulting profile will be a mix of effects of loss of QK5, 6, and 7. Overall, we see clear effects of QKI mutation on ribosome occupancy with 241 nominally upregulated and 115 nominally downregulated genes(**Table S3, Fig. 4E,F**). Consistent with TRAP sensitively assessing the mutant cells, these effects were largely not apparent in the Input RNAseq, demonstrating the need for the targeted enrichment(**Fig. 4G**). Examining the regulated transcripts, in spite of this being a mix of direct and indirect effects, a small but significant fraction of 3’ UTR CLIP targets of QKI-6 are significantly modulated(OR=5.07, FET p<.0018), especially those found in astrocytes(OR=8.8 FET p<.0016). Both up and down regulated transcripts had significant overlap with CLIP targets and those previously described as being translated in PAPs (**Fig. 4H)**, with the KO losing ribosome occupancy of transcripts from PAPs. Looking at ribosome occupancy across all annotated CLIP targets, there was no obvious consistent change, though many of those genes are not expressed in astrocytes. Therefore, subsetting to only CLIP targets expressed in astrocytes, we saw that transcripts from QKI-mutants exhibited a small but significant decrease in ribosome occupancy(average 10% decrease, p<2E-12, T-test)**(Fig. 4I,J)**. These data are consistent with a role for QKI normally in increasing either transcript stability and/or ribosome occupancy of the majority of its direct targets in astrocytes, and is of a magnitude similar to what was previously seen in HEK cells^34^. CLIP targets bound in 5’ UTR, though rare(n=31), had a similar effect(15% decrease, p<9.1E-5, not shown). In contrast, rare targets with QKI-6 binding in introns did not show a consistent direction of effect. It is worth noting the resultant impact on the final protein and mRNA levels following QKI deletion could be mixed as QKI may have distinct roles across its varied targets, or that deletion of all nuclear and cytoplasmic QKI isoforms across postnatal development results in a complicated mix of splicing, translational, and/other indirect effects within the cells. For example *Pbk* and *Ube2c* genes do not appear to be QKI-6 CLIP targets, but are upregulated on QKI loss(**Fig. 4F)**.

As our gRNA targets all isoforms, QKI deletion could also feasibly impact splicing. In a transcript isoform level analysis, we detected 949 transcripts with differential expression arising from 833 genes, including all of those found with the gene level of analysis. Though not our focus here, we have provided these results as a supplemental table for those interested in potential splicing roles(Table S4). For the 74 genes with more than one isoform changing, 40 had isoforms changing in reciprocal directions, which could plausibly represent targets of alternative splicing by QKI-5.

Regardless, analysis of transcripts altering abundance on ribosomes can indicate the function of QKI in GFAP positive cells during postnatal astroglial maturation. Conducting a pathway analysis on the genes upregulated in CRISPR-TRAPseq by loss of QKI indicated a role in cell cycle, especially M-phase(1.4E-8, corrected p-value), and a variety of mitochondrial proteins involved in electron transport chain(4.1E-7)(**Fig. 5A, Table S5)**. The relation to cell cycle in particular could be consistent with a role for QKI in astrocyte maturation, similar to what was previously reported for oligodendrocyte maturation^17^. For example, the mitosis activated kinase Pbk^37^ is upregulated with QKI mutation. Thus, QKI may be required for decreasing proliferation and increasing fate choice, as GFAP positive progenitors from P1 have been shown to contribute to both astrocyte and oligodendroglial lineage^38,39^. To first test this hypothesis, we used CSEA analysis of CRISPR-TRAPseq targets(**Fig. 5B,C**). We saw that QKI mutant astrocytes showed more expression of genes associated with ‘OPC’ cells, a proliferating glial progenitor with potential to turn into both mature astrocytes and oligodendroctyes^40^. Prior work has shown QKI knockout in embryonic neural stem cells prevented their differentiation^28^. Thus, QKI mutation in postnatal GFAP+ cells may influence maturation of astrocytes as well. To further test this hypothesis, we generated a list of genes enriched by TRAP in immature(P7) vs. mature(P32) astrocytes^15^and compared them to transcripts altered by QKI mutation. We found that indeed, deleting QKI resulted in prolonged P7 gene expression and decreased P32 gene expression(**Fig. 5D,E**), consistent with QKI being necessary for astrocyte maturation.

**Figure 5:**
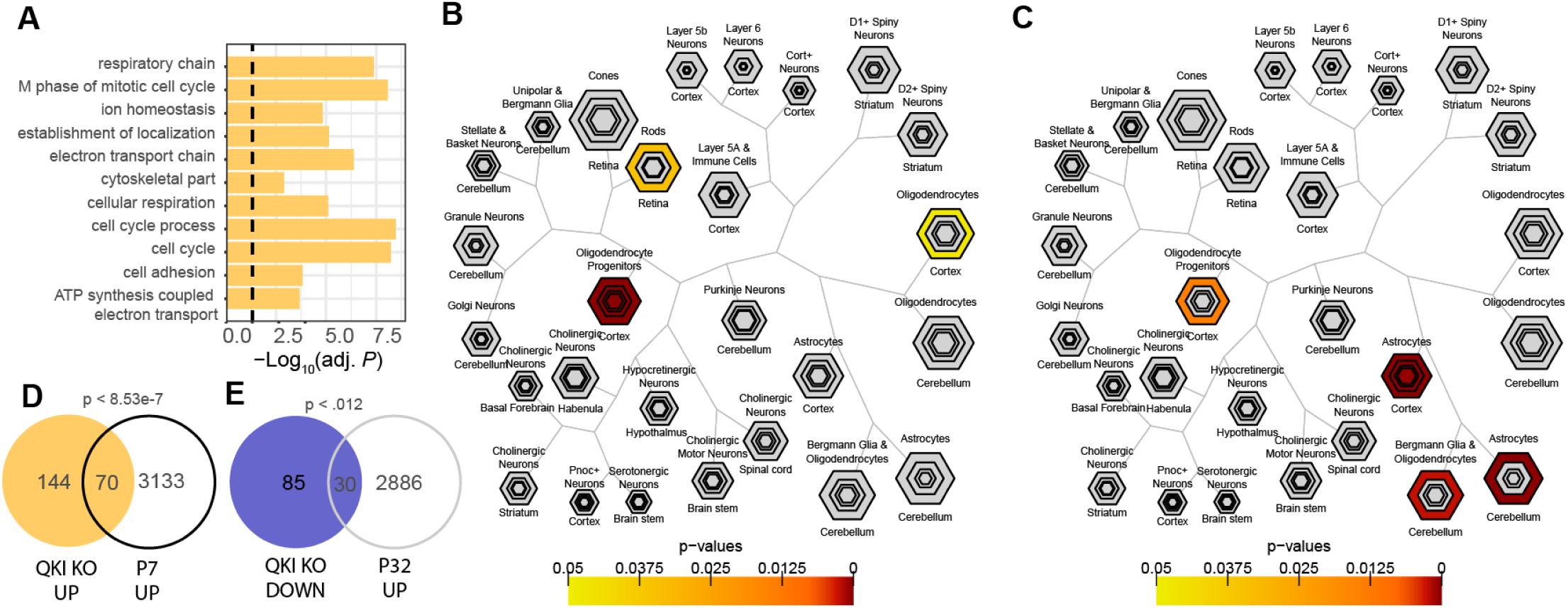
QKI deletion delays astrocyte transcriptional maturation. **A)** Gene Ontology analysis of biological processes enriched by QKI loss reveal increased mitochondrial and cell proliferation. **B)** CSEA of transcripts increased when QKI is deleted are disproportionately found in oligodendroglial progenitor cells. **C)** CSEA of transcripts decreased when QKI is deleted are disproportionately found in adult astrocytes. **D)** Transcripts increased when QKI deleted contain proportionally more P7 astrocyte transcripts. **E)** Transcripts decreased when QKI deleted contain proportionally more P32 astrocyte transcripts.

Overall, our data are consistent with QKI deletion resulting in a mix of direct and indirect cell-autonomous effects, with contributions of all QKI isoforms, and a final consequence on suppressing astroglial maturation.

### Loss of QKI alters local control of synapse numbers and enhancers translation of Sparc mRNA

We next asked whether loss of QKI has functional consequences for its targets. Astrocytes are known controllers of synaptogenesis^41,42^ and astrocyte maturation and synaptic maturation are intimately entwined^43^. One key example is formation of thalamocortical synapses where astrocyte proteins can both promote(via HEVIN) and antagonize(via SPARC) development of this subclass of excitatory synapses^44^. As *Sparc* mRNA is a QKI CLIP target, we examined the abundance of molecularly-defined synapses in the territories of QKI-deleted astrocytes compared to transduced but Cas9-controls. To accomplish this, we stained P21 mouse brains for the presynaptic marker VGLUT2 and post-synaptic marker PSD95, and quantified their apposition in layers IV-V of the cortex where thalamocortical terminals are found^43,45^. We found a significant increase in number of apposed VGLUT2+ and PSD95+ terminals within each astrocyte’s territory in Cas9+ mice, compared to Cas9– littermates(**Fig. 6A,B**), suggesting QKI-mediated regulation of its targets is important in control of synapse numbers. Moreover, this increase in synapses was accompanied by a corresponding increase individually in PSD95 and VGLUT2, within QKI-null astrocyte territories(**Fig S5A,B**). Further, these findings cannot simply be explained by larger astrocytes in Cas9+ mice, as astrocyte area was not significantly different between genotypes(**Fig S5C**). Taken together, these data indicate that astrocytic QKI regulates thalamocortical synapse numbers.

**Figure 6:**
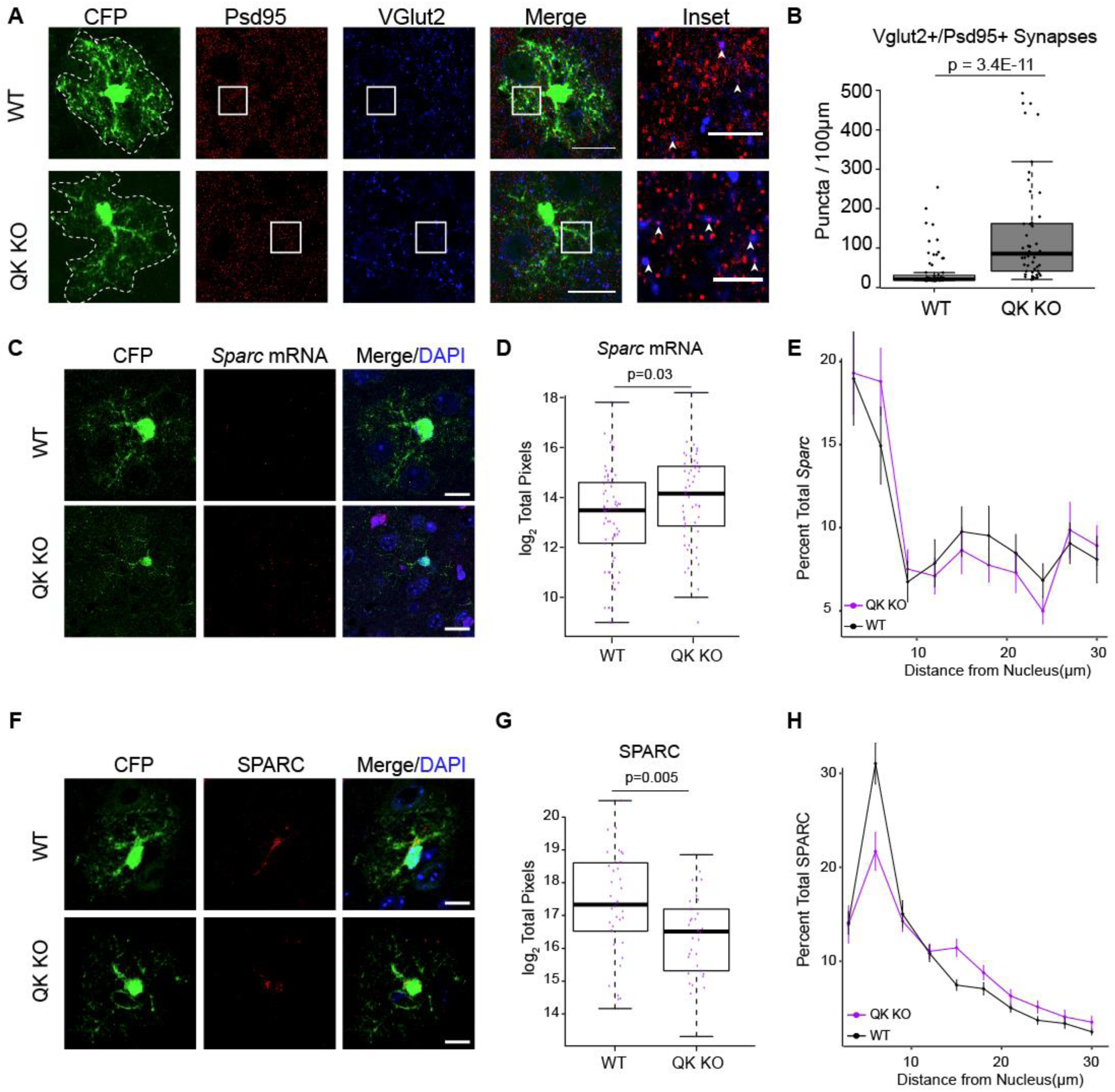
Astrocyte QKI is required for normal excitatory synapse numbers in thalamocortical target layers. **A)** Representative images of PSD95(red), VGLUT2(blue) IF in QKI knockdown astrocytes. **B)** Quantification of colocalized PSD95/VGLUT2 synapses within CFP+ astrocyte territories. N(cells)=51(Cas9+), 67(Cas9-). P values represent Wilcoxon-Rank Sum tests. **C)** Representative images of *Sparc* FISH(red) in QKI knockdown mice. Scale bar = 10μm. D**E)** Total *Sparc* FISH pixels reveals derepression of *Sparc* in Cas9+ cells. P value represents results from Student’s t-test. **E)** Quantification of *Sparc* pixels by distance from the nucleus reveals no significant effects of QKI on *Sparc* mRNA localization. N(Cas9+)=54, N(Cas9-)=66. Repeated measures ANOVA F[genotype]_(1,118)_=2.573,p = 0.111; F[distance]_(1,1078)_=46.467, p < 1.55E-11; F[distance:genotype]_(1,1078)_=0.507, p = 0.476. **F)** Representative images of SPARC IF in QKI knockdown mice. Scale bar = 10μm. **G)** Quantification of total SPARC pixels reveals significantly less SPARC immunoreactivity with loss of QK. P value represents results from Student’s t-test. N(Cas9+)=34, N(Cas9-)=38. **H)** Quantification of percent of SPARC by distance from the nucleus. Repeated measures ANOVA F[genotype]_(1,715)_=0.00,p = 0.9934; F[distance]_(1,715)_=418.581, p < 2.2E-16; F[distance:genotype]_(1,715)_=8.376, p = 0.004.

As SPARC has a known role in regulating these synapse numbers^44^, we hypothesized that perhaps there is reduced SPARC protein produced by these cells. We thus probed abundance, localization or translational efficiency of *Sparc* mRNA and protein by IF. Quantification of *Sparc* mRNA pixels revealed a significant increase in Cas9+ astrocytes, compared to control cells(**Fig. 6C,D**). As RBPs can have more than one function on a target, we also tested if QKI altered mRNA localization by quantifying the abundance of *Sparc* mRNA at increasing distances from the nucleus, and whether or not it was retained in the nucleus as MBP mRNA in QK^v^ mutants. No significant increase in nuclear mRNA was observed between genotypes, and further no main effect of genotype on mRNA was observed(**Fig. 6E**). However, when we quantified total endogenous SPARC protein using the same method we found significantly less SPARC protein in Cas9+ cell territories(**Fig. 6F,G**). Taken together, the concomitant increase in *Sparc* mRNA and decrease in SPARC suggest a role of QKI in regulating translational efficiency of this transcript. Likewise, when we investigated the distribution of SPARC throughout the astrocyte after QKI knockdown and normalized for the change in total abundance, we found that at further distances from the nucleus there was a trend for increased SPARC with a corresponding decrease in the proximal cytoplasm(**Fig. 6H**). Collectively, these data support a role for QKI in regulating proper protein localization and abundance, perhaps by controlling translational efficiency and stability of *Sparc* mRNA. These data are consistent with SPARC dysregulation playing a role in synapse number alteration in the territories of mutant astrocytes.

## Discussion

Here, we define targets of QKI-6 *in vivo* using CLIPseq on mouse forebrain. We initially focused on what QKI-6’s binding pattern indicates about function. First, these data reveal, similar to *in vitro* studies^32^, that QKI-6 target peaks generally reside in 3’ UTRs, supporting a role of QKI in translational control^46^ and mRNA stability^47^. Consistent with binding having important biological roles, bound regions were more conserved and contained enrichment of QBMs. Second, peaks were enriched near stop codons and PolyA signals, and while enriched in QBMs, many sites did not contain them. Further binding was elevated even hundreds of nucleotides from the QBM/peak of highest binding. As QKI is known to homodimerize^11,34^, this spreading binding could be consistent with a cooperativity model where a high-affinity site recruits QKI, which then facilitates additional binding at lower-affinity sites nearby, either through the additional domains in the same molecule, or across dimerized QKI complexes. Of particular interest is the pattern of dual peaks flanking stop codons**(Fig 2F).** As QKI homodimers have been thought to encourage looping of RNA^34,48^, such looping across the stop codon could have important consequences for translational elongation or termination, and could be of interest for future studies.

Motivated by curiosity about translational regulation in astrocytes, we next focused on the significant subset of CLIP targets known to be expressed in them. Our data showed an enrichment of QKI-6 targets in PAP-enriched transcripts^5^, consistent with QKI-6 having some role in regulation of ribosome occupancy in astrocytes. Perhaps similar to its documented role on MBP mRNA, this could be related to nuclear export or pausing during transport, rather than mediating localization per se^20^. QKI-6-bound transcripts include regulators of synapse numbers(*Sparc, Sparcl1*), neurotransmitter uptake(*Slc1a2, Slc1a3*), and astrocytic cytoskeleton(*Gfap*). Interestingly, *Gfap* is a known QKI target in human astrocytes^25^, and other cytoskeletal genes are QKI-regulated in oligodendrocytes^18^, leading to a hypothesis that QKI controls astrocyte development and/or astrogliosis in injury and disease.

Testing this idea however, represented a challenge. The most elegant approaches for defining gene function are deletion studies. However, astrocyte maturation and synaptogenesis are *in vivo* processes occurring over the first weeks of postnatal life. Yet, germline deletion of QKI is embryonic lethal^49^. While the Cre-Lox system is sometimes used to circumvent these issues, during the course of these studies Floxed QKI mice were not available. Furthermore, even then early postnatal studies remain challenging – constitutive astrocyte Cre lines are also expressed in radial glia in embryonic brain^39,50^, where QKI has clear roles in neural stem cells^28^. It is likely brain development would be substantially perturbed obscuring any of the later cell-autonomous effects in maturing astroglia. Some inducible Cre lines for astrocytes exist^51^, however, perinatal tamoxifen injection both dramatically alters sexual differentiation – an effect previously reported in rats that we replicated in mouse (*not shown) -* and leads to substantial rates of delayed lethality: we saw 43% mortality before weaning across 57 pups injected with >100 ug of tamoxifen before P7. Furthermore, an ideal approach would perturb relatively few cells to limit non-cell-autonomous effects on development overall, and would be coupled to a method to specifically yet comprehensively profile cell-autonomous effects. To this end, we developed CRISPR-TRAPseq, wherein sparse gene deletion in specific cell-types, mediated by postnatal gene editing, is coupled to TRAP for specific profiling of targeted cells. We utilized this method to define the consequences of QKI mutation on transcriptome-wide ribosome occupancy in astrocyte development. We saw alterations due to both changes in splicing as well as transcript levels on ribosomes, and these consequences included both direct(CLIP-target) and indirect effects within astrocytes. Generally, there was a tendency for QK loss to decrease ribosome occupancy of QK-6 CLIP targets, perhaps reflecting a loss of transcript stability as seen in cell line QK mutant studies.^34^ This result could be consistent with QK stabilizing these mRNAs, increasing translation initiation, or perhaps stalling mRNAs on ribosomes. Nonetheless, gestalt analysis across altered transcripts argues that QKI is required for proper astrocyte maturation.

The CRISPR-TRAPseq approach proved useful in systematically defining the consequences of QKI deletion in sparse astroglia in a normally developing brain, and we believe it similarly useful for other contexts. For example, swapping promoters will allow targeting of different cell-types, as could electroporation at defined timepoints, provided yields were sufficient. Likewise, gRNAs could target any gene, though RBPs and transcription factors will be especially amenable to TRAP-based readouts. It was also encouraging that TRAP RNA appeared similar in purity with standard bacTRAP(**Fig S4**). Thus, CRISPR-TRAP should be an efficient means to define gene functions *in vivo*. It is important to note that not all cells showed deletion. We suspect remaining cells had silent or missense mutations that disrupted further gRNA binding without altering QKI expression. Adding multiple exon-targeting gRNAs to each vector should remove this vestigial expression. Also, for this study, we delivered Cas9 and TRAP via Rosa alleles and this allowed for a well-controlled littermate design but this design can be implemented via similar approaches delivering these components exogenously(e.g. electroporation or AAV) enabling access to other species. Likewise, existing TRAP and Cas9 FLEX cassettes would enable access to the catalog of available Cre mice, further broadening applicability of the approach. Here, focusing on astrocyte translation, we used this approach coupled to a comprehensive CLIP analysis to describe a key role for QKI in astrocyte maturation and from thence synaptogenesis: As maturation of astrocytes is important for their neuronal interactions,^43,52^ a further consequence was dysregulated synaptogenesis specifically within the QKI mutant astrocyte’s territory. In considering possible mechanisms, we focused on *Sparc*, which controls thalamocortical synapse numbers, and changes expression across age. Previously using 3’ UTR reporter constructs^5^, and here using mosaic QKI mutation, we found that QKI controls translational efficiency of *Sparc* and ultimately alters the protein within the astrocyte domain. This is consistent again with QKI preventing expression during transport of *Sparc* bound for PAPs. It would be interesting to know if this is a recurrent role, or if QKI’s function varies across transcripts. Nonetheless, this study lays groundwork for future studies of additional QKI targets and their roles in astrocyte development and function.

## Material and Methods

### Mice

All procedures were performed in accordance with the guidelines of Washington University’s Institutional Animal Care and Use Committee. Mice were maintained in standard housing conditions with food and water provided *ad libitum*, and crossed at each generation to wildtype C57BL/6J mice from Jackson labs.

### Generation of Cas9 Rosa mice

To generate a Cre-dependent Cas9-expressing mouse line, we engineered Cas9 into a targeting vector containing Rosa26 homology arms (∼1 kb, PCR’s from genome), flanking a CAG enhancer, Lox Stop Lox cassette (adapted from the Allen AI9 constructs, Addgene Plasmid #22799), with 3xFlag-NLS-Cas9-NLS cloned downstream of the stop cassette. The Flag tag is in frame to facilitate confirmation of protein expression. Targeting vector was sequence confirmed then purified for injection into C57BL/6 X CBA F1 hybrid oocytes in conjunction with mRNA coding for a pair of custom TALENs designed to target the mouse ROSA26 locus between the homology arms using the ZiFit targeter (http://zifit.partners.org) and ROSA26 TALENs binding sites as follows:

1. 5’ ROSA26 TALEN binding site: 5’ tccctcgtgatctgcaactcc 3’
2. 3’ ROSA26 TALEN binding site: 5’ gggcgggagtcttctgggca 3’

The TALEN kit used for TALE assembly was a gift from Keith Joung (Addgene kit # 1000000017). DNA fragments encoding ROSA TALEN repeat arrays were cloned into plasmid pJDS71. ROSA TALENs plasmids were linearized for *in vitro* transcription with EcoRI and TALENs RNA was synthesized using the mMessage mMachine T7 Ultra kit (Ambion) and purified with Megaclear columns (Ambion). The cassette of interest was introduced into the mouse genome via pronuclear injection of in vitro transcribed TALENs RNA and ROSA donor DNA. Founders with correctly targeting mutant alleles were identified using long-range PCR with ROSA specific oligos outside of the homology arms and insert specific oligos. Founders may be mosaic with multiple mutant alleles and thus are bred to WT to generate heterozygous F1 offspring. Analysis of F1 offspring via long-range PCR confirmed germline transmission of the correctly targeted allele.

Such progeny of founder MR04 were then bred to a ubiquitous germline Actin-Cre mouse line to confirm Cre-dependence of Cas9 expression (Figure S3). Subsequent generations of Cre-mice were backcrossed to C57BL/6J wildtype mice.

Cas9 efficacy in gRNA-mediated gene targeting was confirmed via loss of QK protein subsequent to delivery of QKI targeting guide RNAs in Cre-expressing transduced astrocytes in vivo (Figure 4A,B).

### CLIP

I. Dissection, lysis and RNAse treatment: Five P21 C57BL/6J mice were anesthetized by inhalation of isoflurane. Mice were decapitated and brains were quickly dissected and moved to ice-cold PBS. The forebrain was removed and dropped into liquid nitrogen in a ceramic mortar and then powdered with a ceramic pestle. Powdered brains were kept in 6cm petri dishes on dry ice until cross-linking. UV cross-linking was performed in a Stratalinker, at 400mJ/cm^2^, 3 times with agitation of the powder between runs to ensure even cross-linking. One sample was not cross-linked, as a control(No XL). Powdered brains were then resuspended in 1mL ice cold 1X Lysis Buffer(50mM Tris-HCl pH 7.4, 100mM NaCl, 1%NP-40, 1X Roche c0mplete EDTA-free protease inhibitor, 1U/uL recombinant RNAsin(Promega), 10mM Na_3_VO_4_, 10mM NaF) and homogenized with a drill(power 14) 10 times. Samples were incubated on ice for 5 minutes then immediately treated with appropriate concentration of RNAse I_f_(NEB) for 3 mins at 37C at 1200RPM. Samples were centrifuged at 20,000xg at 4C for 20 minutes. 2% of the supernatant was kept for Input sample, the rest of the superantant was kept for immunoprecipitation.
II. Immunoprecipitation: Per IP, 6ul of QKI-6 antibody(Millipore #AB9906) or total Rabbit IgG(Jackson Immunoresearch #711-005-152) was added to 72ul of M280 Streptavidin Dynabeads(Invitrogen) and 10ug of Biotinylated Protein G and incubated for 1 hour at room temperature with rotation. Beads were subsequently washed 5 times with 0.1% BSA(Jackson Immunoresearch) before mixed with supernatant. Immunoprecipitation proceeded for 2 hours at 4C with rotation, after which beads were washed with 1X High Salt Wash Buffer(50mM Tris-HCl pH 7.4, 350mM NaCl, 1%NP-40 and 1 unit/ul Promega recombinant RNAsin). In the last wash, beads were split for RNA extraction(98%) and western blot(2%).
III. RNA extraction: Beads were resuspended in 1X LDS Bolt Non-Reducing Sample Buffer, heated for 10 min at 70C then run out on a 4-12% Novex NuPAGE gel in 1X NuPAGE MOPS buffer with a protein ladder(Bio-Rad). The gel was transferred to a 0.2um PVDF membrane for 2 hours at 200mA at 4C. Individual lanes were cut with a razorblade between 37 and 50kDa, to obtain, on average, approximately 100 base length fragments. Membrane strips for each sample were digested in 200ul of 100mM Tris-HCl pH 7.4, 50mM NaCl, 10mM EDTA, 1% Triton-X 100 and 32 units Proteinase K(NEB) at 37C for 1 hour, with shaking. 200ul of 7M Urea was added to each sample for an additional 20minutes at 37C, with shaking. After, 400ul of Acid-Phenol:Chloroform(pH 6.5) was added to each sample, incubated for 5 mins then spun at 10,000xg for 7 minutes at room temperature. The aqueous layer was kept and subsequently cleaned up on the Zymo RNA Clean and Concentrator 5. Quality and quantity of RNA were assessed on an Agilent TapeStation.
IV. Sequencing Library Preparation: Because yields of RNA after CLIP are extremely low(picograms), all RNA harvested was used for library prep. We used direct ligation of an RNA adapter to the harvested RNA after initial dephosphorylation(NEB Antartic Phosphatase) of free ends. RNA was purified using MyONE Silane Dynabeads(Thermo Fisher). Direct ligation of the A01m adapter: (/5Phos/rArGrArUrCrGrGrArArGrArGrCrGrUrCrGrUrGrUrArG/3SpC3/) was added at a final concentration of 1μM using T4 RNA ligase(Enzymatics). Reverse transcription of purified A01m-RNA was carried out using the AR17 primer(ACACGACGCTCTTCCGA) at a final concentration of 1μM, using SuperScript RT III First Strand Synthesis System(Thermo Fisher). RNA and remaining primers were subsequently destroyed with ExoSAP-It(Affymetrix), heat and NaOH. Lastly, a final ligation to cDNA was done to add unique molecular identifiers(allowing removal of PCR duplicates computationally). The Rand103tr3 Adapter: (/5Phos/NNNNNNNNNNAGATCGGAAGAGCACACGTCTG/3SpC3/; Ns indicate any nucleotide) was added at a final concentration of 2μM, using T4 RNA ligase(Enzymatics). PCR(20 total cycles) was then carried out using the Q5 Ultra II Q5 Master Mix(NEB) to amplify final libraries using universal Illumina primers, with sample indexing primers, for sequencing. Detailed protocol is available from the Dougherty Lab upon request.
V. Peak Calling and Differential Expression: Peaks were called across the genome using Piranha^53^ v1.2.1. The mouse genome(mm10) was binned into 50 nucleotide non-overlapping windows. Peaks were called under the genome windows on a merged .bam file from all QKI IP samples. Significant peaks were determined by genome wide FDR p value < 0.05. Count analysis was performed using Subread^54^ v1.5.3, for each sample file. Differential expression was performed using the edgeR package in R. Per edgeR instructions, only peaks with a minimum of five counts(CPM of 41) in two or more samples were analyzed. High confidence QKI peaks were identified by 2-fold enrichment over both IgG IP and Input samples, with an FDR-corrected p value of < 0.1 for discovery purposes. A control set of Input ‘peaks’ is defined as all detected RNAs in the Input matching the CPM cut off described above.

Data are available at GEO: accession pending.

### CLIP analysis

I. MEME Analysis: MEME-CHiP v4.12.0 was performed is discriminative mode, using Input peak sequences as background. Default settings were applied with the exception of using 4 nucleotides as the minimum motif width.
II. Cell-Type Specific Expression Analysis: Analysis for unique genes from QKI or Input significant peaks was performed using the pSI package^55^. Specificity indices were determined using previously published RNA-seq data^21^. A Chi-squared test was used to determine whether the distribution of cell-types in QKI or Input samples were significantly different.
III. Gene Ontology Analysis: GO analysis was performed in Cytoscape v3.5.1 using the BiNGO plug in. Input peak genes were used as the background list. Significant biological process categories were determined by hypergeometric test with Benjamini-Hochberg correction. For analysis of CLIP targets within astrocytes, CLIP targets were subsetted to those genes enriched in astrocytes(logFC >.5 and FDR >.1 in TRAP vs Input comparison, below), and the background for BiNGO analysis was set to all genes enriched in astrocytes.
IV. Analysis of sequence features of targets CLIP Analysis: 25000 transcripts which have 3’ UTR and 3’ UTR length >= 50nt were randomly selected from all mouse (GRCm38.97) transcriptions. The 3’ UTR sequences from a subset of transcripts which have the same length distribution as the CLIP targets were selected as control sequences. The artificial peak location of a control sequence was randomly picked within the 3’ UTR region.

The conservation scores of the CLIP targets and control sequences were retrieved from UCSC basewise conservation scores (phyloP) of 59 vertebrate genomes with mouse. The average conservation scores from CLIP targets are compared with those from control sequences.

Number of polyA sites of each sequence was counted by scanning the polyA signal (A[AT]TAAA). To demonstrate whether the peaks are closer to stop condon or polyA sites, the percentage of distance was calculated as (polyA_site pos – peak pos) / (polyA_site pos – stopcodon pos) * 100 for sequences from plus strand, while (peak pos – polyA site pos) / (stopcodon pos – polyA site pos) * 100 for sequences from minus strand.

Metagene plots were generated by metagene R package 2.14.0 with given alignment files from CLIP, Input and IgG. The expression level of the 1000nt region centered at stop codon, PAS, QBM and peak of CLIP were compared with those of Input and IGG, respectively.

### CRISPR/Cas9 AAV

I. QKI gRNA design and sequences: gRNAs for QKI was designed using the Zhang lab CRISPR design tool(crispr.mit.edu). 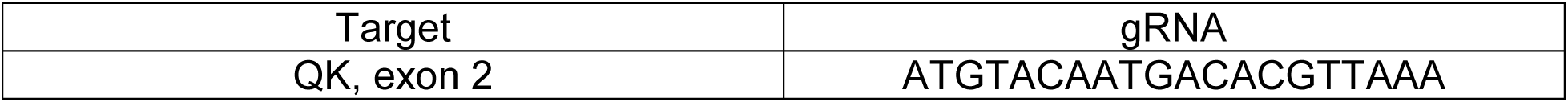
II. QKI knockdown *in vivo*: QKI gRNA was cloned into AAV:ITR-U6-sgRNA(backbone)-pCBh-Cre-WPRE-hGHpA-ITR, a gift from Feng Zhang(Addgene plasmid # 60229). The plasmid was modified such that pCBh promoter was replaced with GFABC1D promoter, and Cre was replaced with CFP and Cre, separated by a 2a peptide. The vector was packaged with AAV9 capsid by the Hope Center for Viral Vectors at Washington University. 1 ul of high titervirus was injected into P0 lox-stop-lox Cas9 knock-in mice(Rosa26:Cas9^tg/+)^ or Cas9-negative littermates (for imaging) or progeny of (Rosa26:Cas9^tg/+^) X (Rosa26:TRAP^tg/tg^) for CRISPR-TRAPseq. Mice were anesthetized on ice for 5 minutes before injections and allowed to recover on heating pad for 15 minutes. Injections were performed as previously described^5^.

### CRISPR-TRAPSeq

Six Cas9+ and six Cas9-negative littermates, all with the TRAP allele, each received intracranial injections of the QKI gRNA AAV (see above) at P1-3 and were sacrificed at p21 for CRISPR/TRAP-Seq. TRAP and PAP-TRAP were performed on p21 cortex homogenates as described previously^5^. RNA quality and concentration were assessed using RNA PicoChips on the Agilent BioAnalyzer following manufacturer’s instructions. Only RNA samples with RIN scores > 7.5 were submitted for library preparation and RNA sequencing.

### RNA Sequencing and Analysis

Library preparation was performed with 0.1-5ng of total RNA, integrity was determined using an Agilent bioanalyzer. ds-cDNA was prepared using the SMARTer Ultra Low RNA kit for Illumina Sequencing (Clontech) per manufacturer’s protocol. cDNA was fragmented using a Covaris E220 sonicator using peak incident power 18, duty factor 20%, cycles/burst 50, time 120 seconds. cDNA was blunt ended, had an A base added to the 3’ ends, and then had Illumina sequencing adapters ligated to the ends. Ligated fragments were then amplified for 14 cycles using primers incorporating unique dual index tags. Fragments were sequenced on an Illumina NovaSeq using paired reads extending 150 bases targeting 30M reads per sample.

Basecalls and demultiplexing were performed with Illumina’s bcl2fastq software and a custom python demultiplexing program with a maximum of one mismatch in the indexing read. Sequencing results were quality checked using FastQC version 0.11.7. Reads aligned to the mouse rRNA were removed by bowtie2 version 2.3.5. Illumina sequencing adaptors were removed and the remaining RNA-seq reads were then aligned to the Ensembl release 97 primary assembly with STAR version 2.5.1a. Gene counts were derived from the number of uniquely aligned unambiguous reads by htseq-count version 0.9.1. Isoform expression of known Ensembl transcripts was estimated with Salmon version 0.8.2. Sequencing performance was assessed for the total number of aligned reads, total number of uniquely aligned reads, and features detected. The ribosomal fraction, known junction saturation, and read distribution over known gene models were quantified with RSeQC version 2.6.2.

For gene level analyses, all gene counts were then imported into the R/Bioconductor package EdgeR5 and TMM normalization size factors were calculated to adjust samples for differences in library size. Only genes that have sufficiently large counts in the smallest group size samples were retained for further analysis using the default settings of the filterByExpr function in EdgeR. Differential expression analysis was then performed to analyze for differences between conditions. A negative binomial generalized log-linear model (GLM) was fit to the counts for each gene. Then the likelihood ratio tests (LRT) were conducted for each comparison. Similar approaches were taken for transcript isoforms, but the differential expression analysis used the R/Bioconductor package Limma with the voomWithQualityWeights function to further account for increased levels of variance at the isoform level. Data are available at GEO: GSE146935.

### Immunohistochemistry

Mice were sacrificed by inhalation of isoflurane and then rapidly perfused with ice-cold 1X PBS, followed by 4% paraformaldehyde. Brains were post-fixed overnight and then cryoprotected in 30% sucrose. Sectioning was performed on a cryostat(Leica). IHC was performed on 14μm slide-mounted coronal brain sections in all cases. Slides were blocked in 5% Normal Donkey Serum(Jackson Immunoresearch) with 0.25% Triton X-100 in PBS for at least 30 minutes at room temperature. Primary antibodies were incubated overnight at 4C. Secondary antibodies(Invitrogen) were incubated at 1:1000 for 90 minutes at room temperature. Quantification of total pixels was performed blind to genotype and/or condition using custom macros for ImageJ(NIH), as previously described^5^. N(cells) is represented in the figure legends. Experiments were performed on at least 3 mice per genotype.

| Antibody | Catalog Number / Company | Dilution |
| --- | --- | --- |
| QK5, Rb | AB9904 / Millipore | 1:500 |
| QKI-6, Rb | AB9906 / Millipore | 1:500 |
| QK7, Rb | AB9908 / Millipore | 1:500 |
| GFP, Ck | GFP-1020 / Aves | 1:1000 |
| SPARC, Gt | AF942 / R&D Systems | 1:40(final 5ug/mL) |
| VGLUT2, Gp | AB2251 / Millipore | 1:2500 |
| PSD-95, Rb | 51-6900 / Invitrogen | 1:250 |

### Synapse assay

40μm free-floating sections were blocked in 20% normal donkey serum in 1X PBS for 1 hour at RT. Primary antibodies were incubated overnight at RT with 50RPM shaking in 10% normal donkey serum, 0.3% Triton X-100 in PBS. Secondary antibodies were incubated at room temperature for 4 hours at 1:200 dilutions. Single-plane 0.9μm optical images were collected on a Zeiss confocal microscope. Quantification of pixels within astrocyte territories(ROIs) was done in ImageJ with custom macros using previously published methods^43,56^. We analyzed only the cells in which QKI was effectively lost in Cas9+ animals as assessed by QKI IF on a cell by cell basis.

### Western blotting

Whole mouse brains were dissected after isoflurane inhalation and rapid decapitation. Brains were homogenized(10mM HEPES pH 7.4, 150mM KCl, 5mM MgCl_2_, 0.5mM DTT, supplemented with 1μl rRNAsin(Promega) and 1 μl Superasin(Ambion) per ml and 1 mini EDTA-free protease inhibitor tablet(Roche) per 10ml) using a glass homogenizer and Teflon pestle with a drill. Lysates were cleared by centrifugation at 2000xg for 10 minutes. Concentration of the supernatant was determined by BCA assay(Thermo Fisher). 50ug of total lysate was run on 4-12% SDS-PAGE gel(Bio-rad) and transferred to 0.2μm PVDF membrane(Bio-rad). Ponceau image was quantified for loading control. Membranes were blocked in 5% Milk in 1X TBS with 0.5% Tween for 3 hours at room temperature. Membranes were incubated in antibody overnight at 4C. Appropriate HRP conjugated secondary antibodies were applied after washes at room temperature for 1 hour. Membranes were incubated in Clarity(Bio-rad) chemiluminescence reagents for 5 minutes and then developed for 1 minute in a myECL Imager(Thermo). N is represented in figure legend.

| Antibody | Catalog Number / Company | Dilution |
| --- | --- | --- |
| QKI-6, Rb | AB9906 / Millipore | 1:1000 |

### QKI-6 RNA Immunoprecipitation

Independent P21 mouse forebrains were dissected in 2mL of 1X Lysis Buffer(as above). Homogenates were spun at 20,000xg for 20minutes and supernatant was collected. 2.5% of the supernatant was saved for western blotting in 1X Laemmli Buffer(Bio-Rad) or RNA extraction. The rest of the lysate was incubated with IgG or QKI-6 coupled streptavidin Dynabeads for 2 hours at 4C(prepared identically to CLIP protocol). Beads were washed in 1X High Salt Wash Buffer, 4 times for 5 minutes each with end-over-end rotation. 10% of the beads were resuspended in 1X Laemmli buffer and loaded onto an SDS-PAGE gel to confirm immunoprecipitation of QK. RNA was extracted from the remaining 90% of beads.

### Fluorescent *in situ* hybridization

FISH was performed as described previously, using a previously published probe for *Sparc* ^5^. Briefly, 14μm slide-mounted coronal brain sections were hybridized with 100ng of purified probe overnight at 65C. Signal was amplified using a tyramide signal amplification Cy3 kit(Perkin Elmer). IHC was performed following FISH identically to IHC described above. Quantification of total pixels was performed blind to genotype and condition using custom macros for ImageJ(NIH). N(cells) is represented in the figure legends. Experiments were performed on at least 3 mice per genotype.

## Acknowledgements

This work was supported by 5R01NS102272 and R01MH116999. We thank Dr. Philip Uren for helpful discussions and assistance using Piranha, and Dr. Laura Clarke for P7 and P32 astrocyte data, and Jess Hoisington-Lopez, Scott Lee, Katherine B. McCullough and Stephen Plassmeyer for technical contributions. We thank the Genome Technology Access Center in the Department of Genetics at Washington University School of Medicine for help with genomic analysis. The Center is partially supported by NCI Cancer Center Support Grant #P30 CA91842 to the Siteman Cancer Center and by ICTS/CTSA Grant# UL1TR002345.

## Supplemental Figures

**Figure S1:**
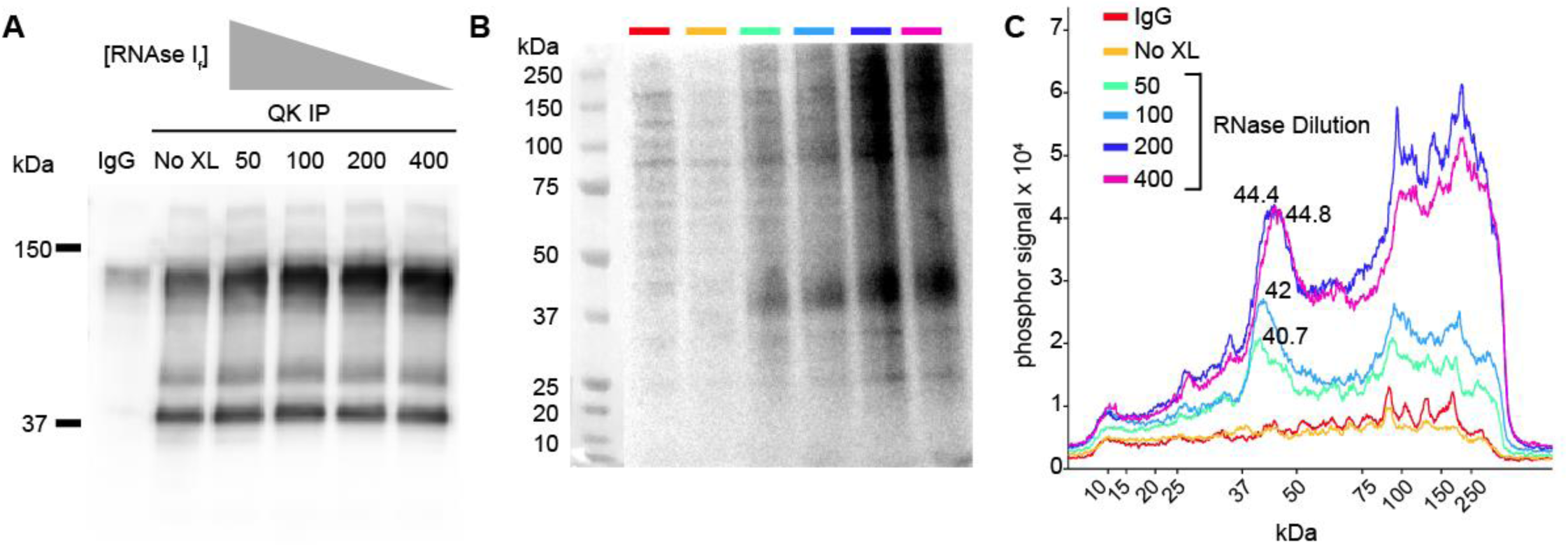
Optimization of QKI CLIP protocol. **A)** Western blot demonstrates that RNAse I_f_ digestion does not affect QKI IP. QKI-specific band approx. 38kDa. Band below 150 kDa is IgG heavy and light chains(samples prepared in non-reducing sample buffer). IgG IP does not pull down QK. Values along lanes represent dilution of RNAse If. **B)** Phosphor screen image of ^32^P-labeled RNA from QKI IP, across RNAse I_f_ dilutions. Colors above lanes correspond to dilutions of RNAse I_f_ in plot in panel C. **C)** Quantification of ^32^P signal from image in panel B. QKI-bound RNA is around 40-44kDA, and this area of the gel was excised for sequencing libraries.

**Figure S2:**
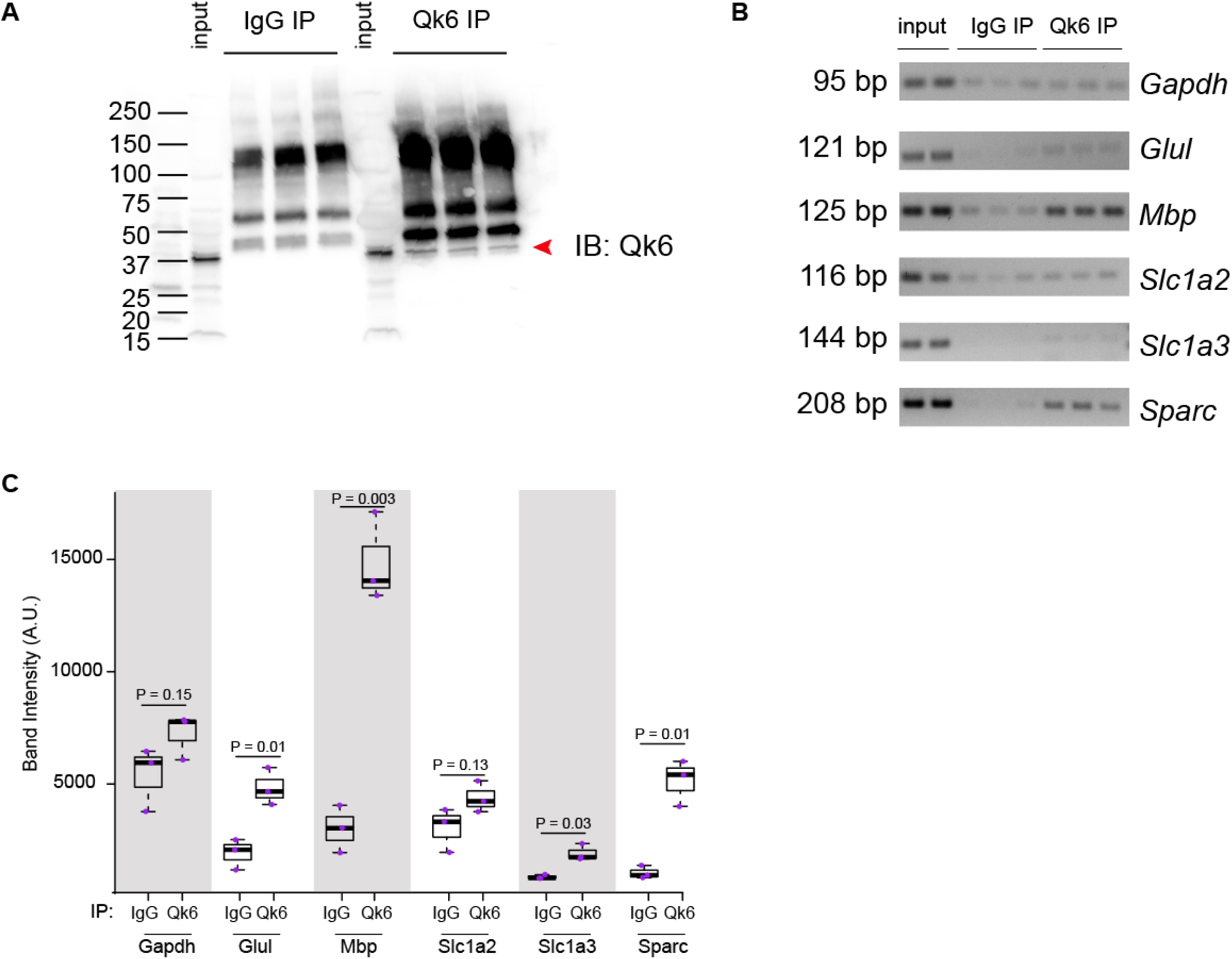
Confirmation of CLIP target enrichment. **A)** Immunoblot of QKI-6 IPs from C57BL/6J mice. IgG IP shows non specific bands. QKI-6-specific band(38kDa) is seen in the Input samples and in all 3 QKI IPs, indicated by the red arrow, but not in the IgG IPs. **B)** RT-PCR gel images. **C)** Densitometric analysis of RT-PCR gel images. Quantification was performed on uncropped, unaltered images. P-values represent Student’s t-tests.

**Figure S3:**
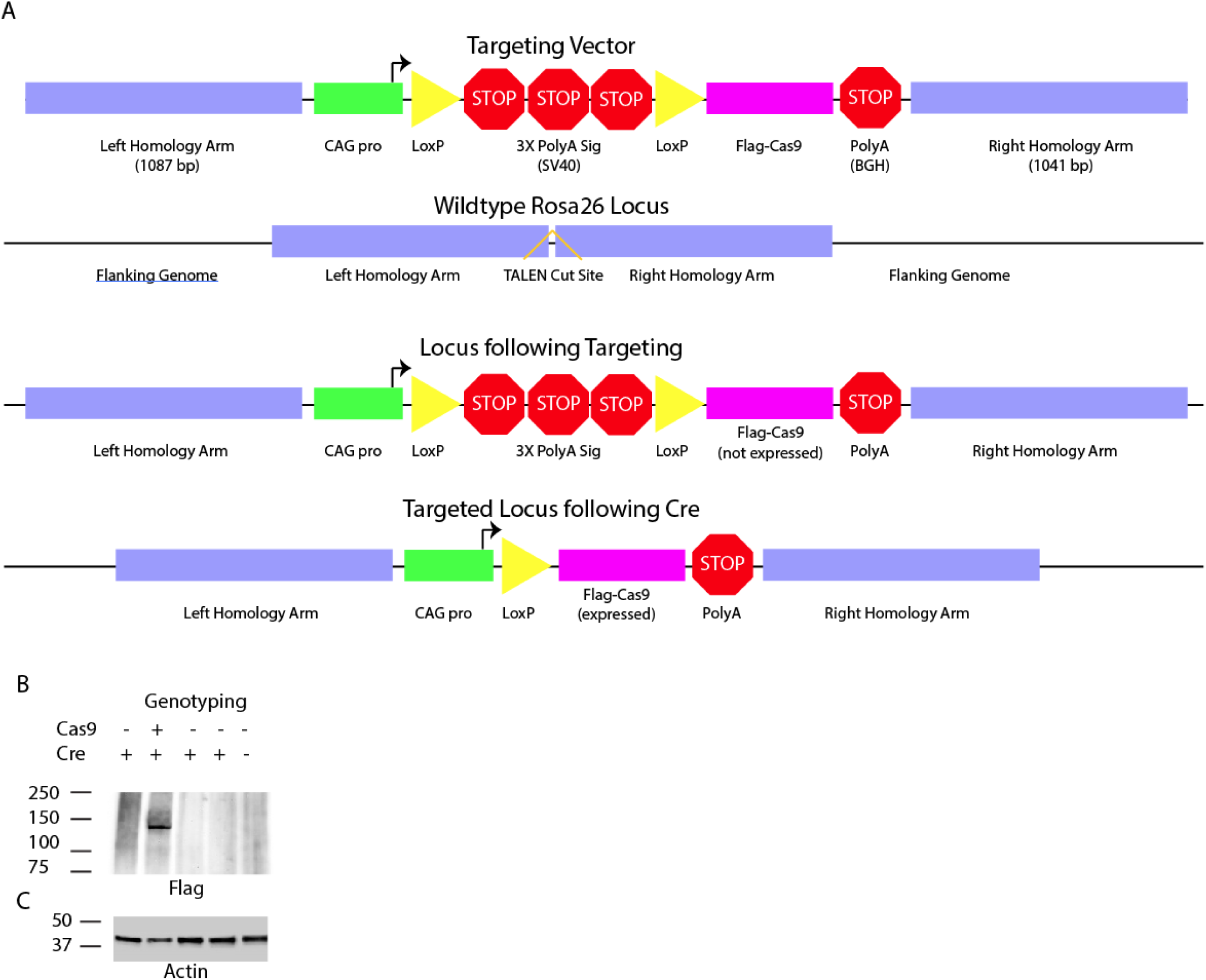
Generation of a Cre dependent Cas9 expressing mouse line. **A)** Schematic of targeting strategy. TALENS targeting the ROSA26 locus were utilized to insert a targeting vector containing a CAG/Rosa hybrid promoter driving expression of Cas9, interrupted by a Lox Stop Loc cassette. Final locus is engineered to express Cas9-Flag in response to Cre recombinase. **B)** Anti-flag immunoblot detects protein only in brains from mice that genotyped positive for both the Cre allele and the Cas9-Flag Rosa allele. **C)** Loading control (anti-actin) showing all lysates contained protein.

**Figure S4:**
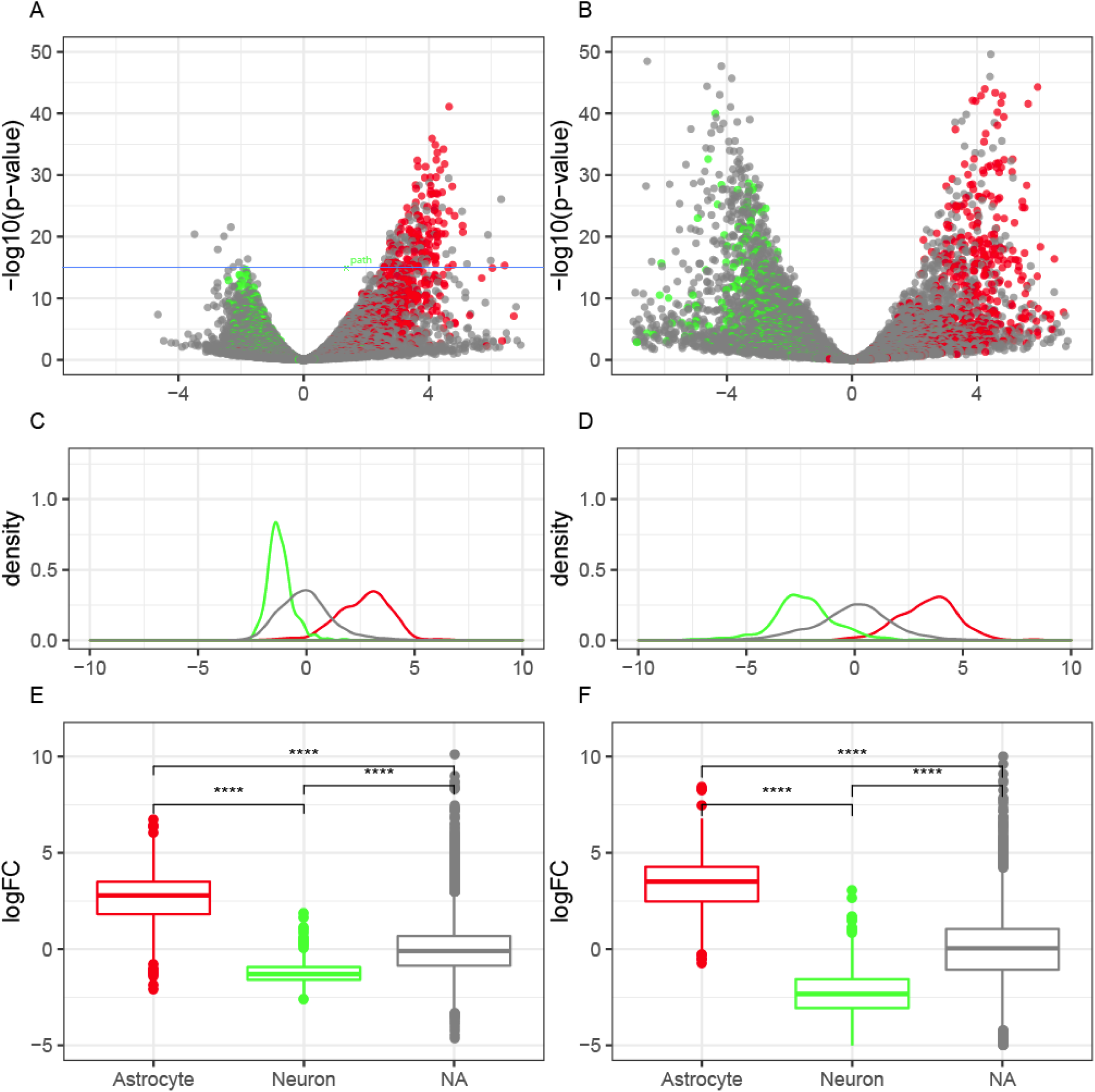
Viral TRAP is comparable to bacTRAP. **A,C,E**) Viral CRISPR-TRAPseq. **B,D,F)** Aldh1l1 bacTRAP. **A,B)** Volcano plots showing enrichment and depletion of known astrocytic (red) and neuronal genes (green). **C,D)** Density of transcripts representing each enrichment. **E,F)** Boxplots showing the fold change of known marker genes (astrocytic, neuronal, or other (NA)).

**Figure S5:**
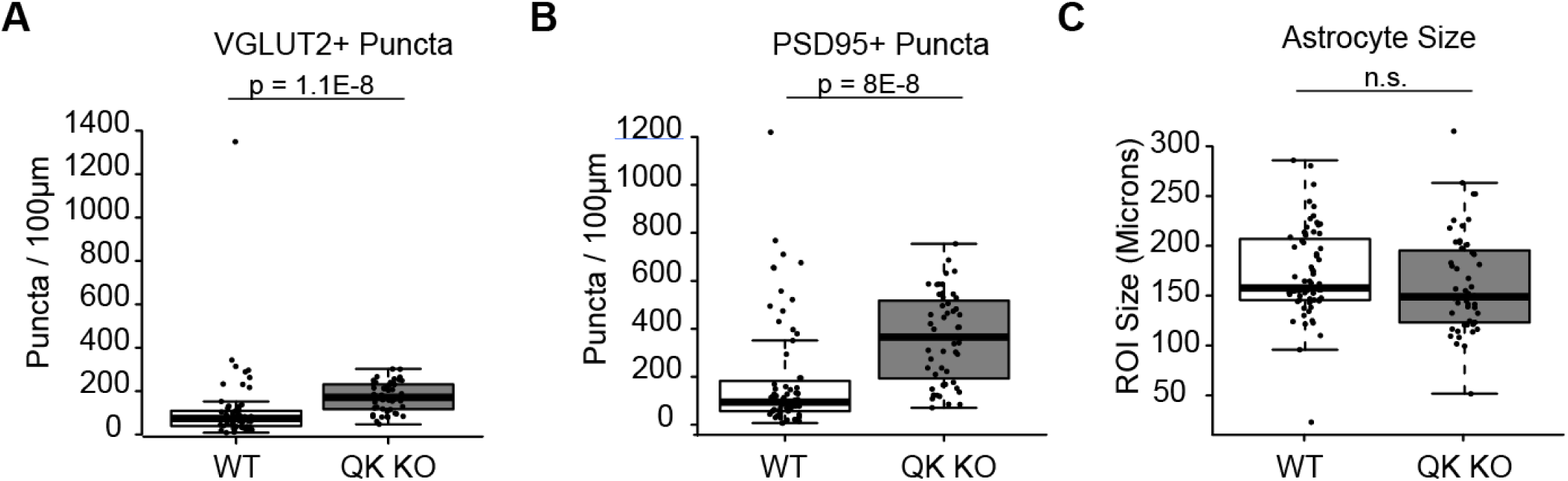
QK KO impacts VGLUT2 and PSD95 expression but not astrocyte size. **A)** Quantification of VGLUT2+ puncta within CFP+ astrocyte territories. **B**) Quantification of Psd9+ puncta within CFP+ astrocyte territories. **C**) Quantification of CFP+ astrocyte area. P values represent Wilcoxon-rank sum test.

## Supplemental Table Legends

**Table S1: QKI-6 bound regions.**

Column headers: logFC, logCPM, LR, PValue, & FDR describe results comparing QKI-6 CLIP enrichment over both Input and IgG controls or EdgeR analysis of candidate QKI-6 bound peaks.

GeneFeature, utr3match, utr5match, intronmatch, cdsmatch, & intergenic describe the gene annotations overlapping with the peak. Sequences, start, & end describe the sequence of the 50bp cpeak, as well as coordinates.

**Table S2: GO analysis of CLIP targets**

BinGO output on all genes bound by QKI-6, reporting any of the full GO categories with an FDR p<.05. Headers as described by BinGO.

**Table S3: Genes with differential ribosome occupancy following QKI deletion in maturing astrocytes.**

EdgeR for all transcripts with p<.05 in this contrast. Headers as above.

**Table S4: Isoform level analysis of differential ribosome occupancy following QKI deletion in maturing astrocytes.**

Limma results for all transcript isoforms with p<.05. Column headers: Ensembl_Transcript_ID and Transcript_Symbol describe the ID of the isoform and the Entrezgene_ID, Ensembl_Gene_ID, Gene Symbol, and Description describe the parent gene of each isoform. Each isoform lists the linear fold-change, log 2 fold-change with confidence intervals, p-value, and Benjamini-Hochberg adjusted p-value.

**Table S5: GO analysis on transcripts altered by QKI deletion.**

BinGO output(FDR p<.005 cutoff) on all transcripts upregulated following QKI deletion. Headers as described by BinGO.

## References

1. Andreassi, C. & Riccio, A. To localize or not to localize: mRNA fate is in 3’UTR ends. Trends Cell Biol. 19, 465–474 (2009).

2. Huang, Y.-S., Carson, J. H., Barbarese, E. & Richter, J. D. Facilitation of dendritic mRNA transport by CPEB. Genes Dev. 17, 638–653 (2003).

3. Darnell, J. C. et al. FMRP stalls ribosomal translocation on mRNAs linked to synaptic function and autism. Cell 146, 247–261 (2011).

4. Wang, E. T. et al. Dysregulation of mRNA Localization and Translation in Genetic Disease. J. Neurosci. Off. J. Soc. Neurosci. 36, 11418–11426 (2016).

5. Sakers, K. et al. Astrocytes locally translate transcripts in their peripheral processes. Proc. Natl. Acad. Sci. U. S. A. 114, E3830–E3838 (2017).

6. Lee, J.-A. et al. Cytoplasmic Rbfox1 Regulates the Expression of Synaptic and Autism-Related Genes. Neuron 89, 113–128 (2016).

7. Boutej, H. et al. Diverging mRNA and Protein Networks in Activated Microglia Reveal SRSF3 Suppresses Translation of Highly Upregulated Innate Immune Transcripts. Cell Rep. 21, 3220–3233 (2017).

8. Li, Z., Zhang, Y., Li, D. & Feng, Y. Destabilization and mislocalization of myelin basic protein mRNAs in quaking dysmyelination lacking the QKI RNA-binding proteins. J. Neurosci. Off. J. Soc. Neurosci. 20, 4944–4953 (2000).

9. Higashimori, H. et al. Selective Deletion of Astroglial FMRP Dysregulates Glutamate Transporter GLT1 and Contributes to Fragile X Syndrome Phenotypes In Vivo. J. Neurosci. Off. J. Soc. Neurosci. 36, 7079–7094 (2016).

10. Pilaz, L.-J., Lennox, A. L., Rouanet, J. P. & Silver, D. L. Dynamic mRNA Transport and Local Translation in Radial Glial Progenitors of the Developing Brain. Curr. Biol. CB 26, 3383–3392 (2016).

11. Chen, T. & Richard, S. Structure-function analysis of Qk1: a lethal point mutation in mouse quaking prevents homodimerization. Mol. Cell. Biol. 18, 4863–4871 (1998).

12. Li, Z. et al. Defective smooth muscle development in qkI-deficient mice. Dev. Growth Differ. 45, 449–462 (2003).

13. Jan, E., Motzny, C. K., Graves, L. E. & Goodwin, E. B. The STAR protein, GLD-1, is a translational regulator of sexual identity in Caenorhabditis elegans. EMBO J. 18, 258–269 (1999).

14. Boisvert, M. M., Erikson, G. A., Shokhirev, M. N. & Allen, N. J. The Aging Astrocyte Transcriptome from Multiple Regions of the Mouse Brain. Cell Rep. 22, 269–285 (2018).

15. Clarke, L. E. et al. Normal aging induces A1-like astrocyte reactivity. Proc. Natl. Acad. Sci. 115, E1896–E1905 (2018).

16. Hardy, R. J. et al. Neural cell type-specific expression of QKI proteins is altered in quakingviable mutant mice. J. Neurosci. Off. J. Soc. Neurosci. 16, 7941–7949 (1996).

17. Larocque, D. et al. Protection of p27(Kip1) mRNA by quaking RNA binding proteins promotes oligodendrocyte differentiation. Nat. Neurosci. 8, 27–33 (2005).

18. Zhao, L. et al. QKI binds MAP1B mRNA and enhances MAP1B expression during oligodendrocyte development. Mol. Biol. Cell 17, 4179–4186 (2006).

19. Doukhanine, E., Gavino, C., Haines, J. D., Almazan, G. & Richard, S. The QKI-6 RNA binding protein regulates actin-interacting protein-1 mRNA stability during oligodendrocyte differentiation. Mol. Biol. Cell 21, 3029–3040 (2010).

20. Larocque, D. et al. Nuclear retention of MBP mRNAs in the quaking viable mice. Neuron 36, 815–829 (2002).

21. Zhang, Y. et al. An RNA-Sequencing Transcriptome and Splicing Database of Glia, Neurons, and Vascular Cells of the Cerebral Cortex. J. Neurosci. 34, 11929–11947 (2014).

22. Zhang, Y. et al. Purification and Characterization of Progenitor and Mature Human Astrocytes Reveals Transcriptional and Functional Differences with Mouse. Neuron 89, 37–53 (2016).

23. Jiang, L., Saetre, P., Radomska, K. J., Jazin, E. & Lindholm Carlström, E. QKI-7 regulates expression of interferon-related genes in human astrocyte glioma cells. PloS One 5, (2010).

24. Wu, J. I., Reed, R. B., Grabowski, P. J. & Artzt, K. Function of quaking in myelination: regulation of alternative splicing. Proc. Natl. Acad. Sci. U. S. A. 99, 4233–4238 (2002).

25. Radomska, K. J. et al. RNA-binding protein QKI regulates Glial fibrillary acidic protein expression in human astrocytes. Hum. Mol. Genet. 22, 1373–1382 (2013).

26. Farnsworth, B. et al. QKI6B mRNA levels are upregulated in schizophrenia and predict GFAP expression. Brain Res. 1669, 63–68 (2017).

27. Ebersole, T. A., Chen, Q., Justice, M. J. & Artzt, K. The quaking gene product necessary in embryogenesis and myelination combines features of RNA binding and signal transduction proteins. Nat. Genet. 12, 260–265 (1996).

28. Hayakawa-Yano, Y. et al. An RNA-binding protein, Qki5, regulates embryonic neural stem cells through pre-mRNA processing in cell adhesion signaling. Genes Dev. 31, 1910–1925 (2017).

29. Bailey, T. L., Johnson, J., Grant, C. E. & Noble, W. S. The MEME Suite. Nucleic Acids Res. 43, W39–49 (2015).

30. Galarneau, A. & Richard, S. Target RNA motif and target mRNAs of the Quaking STAR protein. Nat. Struct. Mol. Biol. 12, 691–698 (2005).

31. Tushev, G. et al. Alternative 3’ UTRs Modify the Localization, Regulatory Potential, Stability, and Plasticity of mRNAs in Neuronal Compartments. Neuron (2018) doi: 10.1016/j.neuron.2018.03.030.

32. Fagg, W. S. et al. Autogenous cross-regulation of Quaking mRNA processing and translation balances Quaking functions in splicing and translation. Genes Dev. 31, 1894–1909 (2017).

33. Theil, K., Herzog, M. & Rajewsky, N. Post-transcriptional Regulation by 3’ UTRs Can Be Masked by Regulatory Elements in 5’ UTRs. Cell Rep. 22, 3217–3226 (2018).

34. Teplova, M. et al. Structure–function studies of STAR family Quaking proteins bound to their in vivo RNA target sites. Genes Dev. 27, 928–940 (2013).

35. Xu, X., Wells, A. B., O’Brien, D. R., Nehorai, A. & Dougherty, J. D. Cell type-specific expression analysis to identify putative cellular mechanisms for neurogenetic disorders. J. Neurosci. Off. J. Soc. Neurosci. 34, 1420–1431 (2014).

36. Wu, H. Y., Dawson, M. R. L., Reynolds, R. & Hardy, R. J. Expression of QKI Proteins and MAP1B Identifies Actively Myelinating Oligodendrocytes in Adult Rat Brain. Mol. Cell. Neurosci. 17, 292–302 (2001).

37. Dougherty, J. D. et al. PBK/TOPK, a Proliferating Neural Progenitor-Specific Mitogen-Activated Protein Kinase Kinase. J. Neurosci. 25, 10773–10785 (2005).

38. Menn, B. et al. Origin of Oligodendrocytes in the Subventricular Zone of the Adult Brain. J. Neurosci. 26, 7907–7918 (2006).

39. Casper, K. B. & McCarthy, K. D. GFAP-positive progenitor cells produce neurons and oligodendrocytes throughout the CNS. Mol. Cell. Neurosci. 31, 676–684 (2006).

40. Suzuki, N. et al. Differentiation of Oligodendrocyte Precursor Cells from Sox10 -Venus Mice to Oligodendrocytes and Astrocytes. Sci. Rep. 7, 1–11 (2017).

41. Allen, N. J. Role of glia in developmental synapse formation. Curr. Opin. Neurobiol. 23, 1027–1033 (2013).

42. Baldwin, K. T. & Eroglu, C. Molecular mechanisms of astrocyte-induced synaptogenesis. Curr. Opin. Neurobiol. 45, 113–120 (2017).

43. Stogsdill, J. A. et al. Astrocytic neuroligins control astrocyte morphogenesis and synaptogenesis. Nature 551, 192–197 (2017).

44. Kucukdereli, H. et al. Control of excitatory CNS synaptogenesis by astrocyte-secreted proteins Hevin and SPARC. Proc. Natl. Acad. Sci. U. S. A. 108, E440–449 (2011).

45. Nahmani, M. & Erisir, A. VGluT2 immunochemistry identifies thalamocortical terminals in layer 4 of adult and developing visual cortex. J. Comp. Neurol. 484, 458–473 (2005).

46. Saccomanno, L. et al. The STAR protein QKI-6 is a translational repressor. Proc. Natl. Acad. Sci. U. S. A. 96, 12605–12610 (1999).

47. Zearfoss, N. R., Clingman, C. C., Farley, B. M., McCoig, L. M. & Ryder, S. P. Quaking Regulates Hnrnpa1 Expression through Its 3’ UTR in Oligodendrocyte Precursor Cells. PLOS Genet 7, e1001269 (2011).

48. Conn, S. J. et al. The RNA binding protein quaking regulates formation of circRNAs. Cell 160, 1125–1134 (2015).

49. Justice, M. J. & Bode, V. C. Three ENU-induced alleles of the murine quaking locus are recessive embryonic lethal mutations. Genet. Res. 51, 95–102 (1988).

50. Foo, L. C. & Dougherty, J. D. Aldh1L1 is expressed by postnatal neural stem cells in vivo. Glia 61, 1533–1541 (2013).

51. Srinivasan, R. et al. New Transgenic Mouse Lines for Selectively Targeting Astrocytes and Studying Calcium Signals in Astrocyte Processes In Situ and In Vivo. Neuron 92, 1181–1195 (2016).

52. Holt, L. M. et al. Astrocyte morphogenesis is dependent on BDNF signaling via astrocytic TrkB.T1. eLife 8, e44667 (2019).

53. Uren, P. J. et al. Site identification in high-throughput RNA-protein interaction data. Bioinforma. Oxf. Engl. 28, 3013–3020 (2012).

54. Liao, Y., Smyth, G. K. & Shi, W. The Subread aligner: fast, accurate and scalable read mapping by seed-and-vote. Nucleic Acids Res. 41, e108 (2013).

55. Dougherty, J. D., Schmidt, E. F., Nakajima, M. & Heintz, N. Analytical approaches to RNA profiling data for the identification of genes enriched in specific cells. Nucleic Acids Res. 38, 4218–4230 (2010).

56. Ippolito, D. M. & Eroglu, C. Quantifying Synapses: an Immunocytochemistry-based Assay to Quantify Synapse Number. J. Vis. Exp. JoVE (2010) doi: 10.3791/2270.

57. Dougherty, J. D. et al. Candidate pathways for promoting differentiation or quiescence of oligodendrocyte progenitor-like cells in glioma. Cancer Res. 72, 4856–4868 (2012).

